# Cooperative Gsx2-DNA Binding Requires DNA Bending and a Novel Gsx2 Homeodomain Interface

**DOI:** 10.1101/2023.12.08.570805

**Authors:** Jordan A. Webb, Edward Farrow, Brittany Cain, Zhenyu Yuan, Alexander E. Yarawsky, Emma Schoch, Ellen K. Gagliani, Andrew B. Herr, Brian Gebelein, Rhett A. Kovall

## Abstract

The conserved Gsx homeodomain (HD) transcription factors specify neural cell fates in animals from flies to mammals. Like many HD proteins, Gsx factors bind A/T-rich DNA sequences prompting the question — how do HD factors that bind similar DNA sequences *in vitro* regulate specific target genes *in vivo*? Prior studies revealed that Gsx factors bind DNA both as a monomer on individual A/T-rich sites and as a cooperative homodimer to two sites spaced precisely seven base pairs apart. However, the mechanistic basis for Gsx DNA binding and cooperativity are poorly understood. Here, we used biochemical, biophysical, structural, and modeling approaches to (1) show that Gsx factors are monomers in solution and require DNA for cooperative complex formation; (2) define the affinity and thermodynamic binding parameters of Gsx2/DNA interactions; (3) solve a high-resolution monomer/DNA structure that reveals Gsx2 induces a 20° bend in DNA; (4) identify a Gsx2 protein-protein interface required for cooperative DNA binding; and (5) determine that flexible spacer DNA sequences enhance Gsx2 cooperativity on dimer sites. Altogether, our results provide a mechanistic basis for understanding the protein and DNA structural determinants that underlie cooperative DNA binding by Gsx factors, thereby providing a deeper understanding of HD specificity.

## INTRODUCTION

Homeodomain (HD) proteins are highly conserved eukaryotic transcription factors (TFs) that control extensive developmental networks during embryogenesis, as well as maintain tissue homeostasis and metabolic responses within adult organisms^1–3^. The HD superfamily accounts for 15-30% of all TFs in plants and animals and is defined by an ∼60 amino acid conserved HD that binds DNA^2^. Despite numerous HD family members regulating diverse functions *in vivo*, previous studies have revealed that the majority of HD TFs bind highly similar A/T-rich DNA motifs *in vitro*^4–6^. This finding raises a long-standing question in the field regarding how HD TFs accurately achieve the *in vivo* specificity required for proper gene regulation when they bind nearly identical DNA motifs *in vitro*.

*Gsx* genes encode HD TFs that are conserved from *Drosophila* to humans^2^ (Figure 1A-B). Gsx2, as well as its paralog Gsx1 and the fly ortholog Ind, regulate specific aspects of neural development including dorsal-ventral patterning and neural fate specification. Within the forebrain, Gsx2 sits atop a gene regulatory network required for proper ventral telencephalon development^7–12^. During early embryogenesis, Gsx2 specifies lateral ganglionic eminence (LGE) progenitors and restricts expression of other telencephalic developmental programs, thus promoting proper dorsal-ventral patterning of the telencephalon^9,10,12^. At later embryonic stages, Gsx2 plays a critical role in regulating neural cell fates by promoting the expression of pro-neurogenic genes, such as the TF Ascl1^8–10,12^. In agreement with these studies, Gsx2 mutations in mice show severe neurological defects, specifically loss of basal ganglia neuronal subtypes^7,8,13–16^. More recently, human patient studies identified two *GSX2* variants associated with severe intellectual disability, spastic tetraparesis, and dystonia^17^. Patient MRI studies revealed nearly complete basal ganglia agenesis, consistent with *Gsx2* mouse studies and the role of the basal ganglia in purposeful movement and cognitive responses^17^.

**Figure 1.**
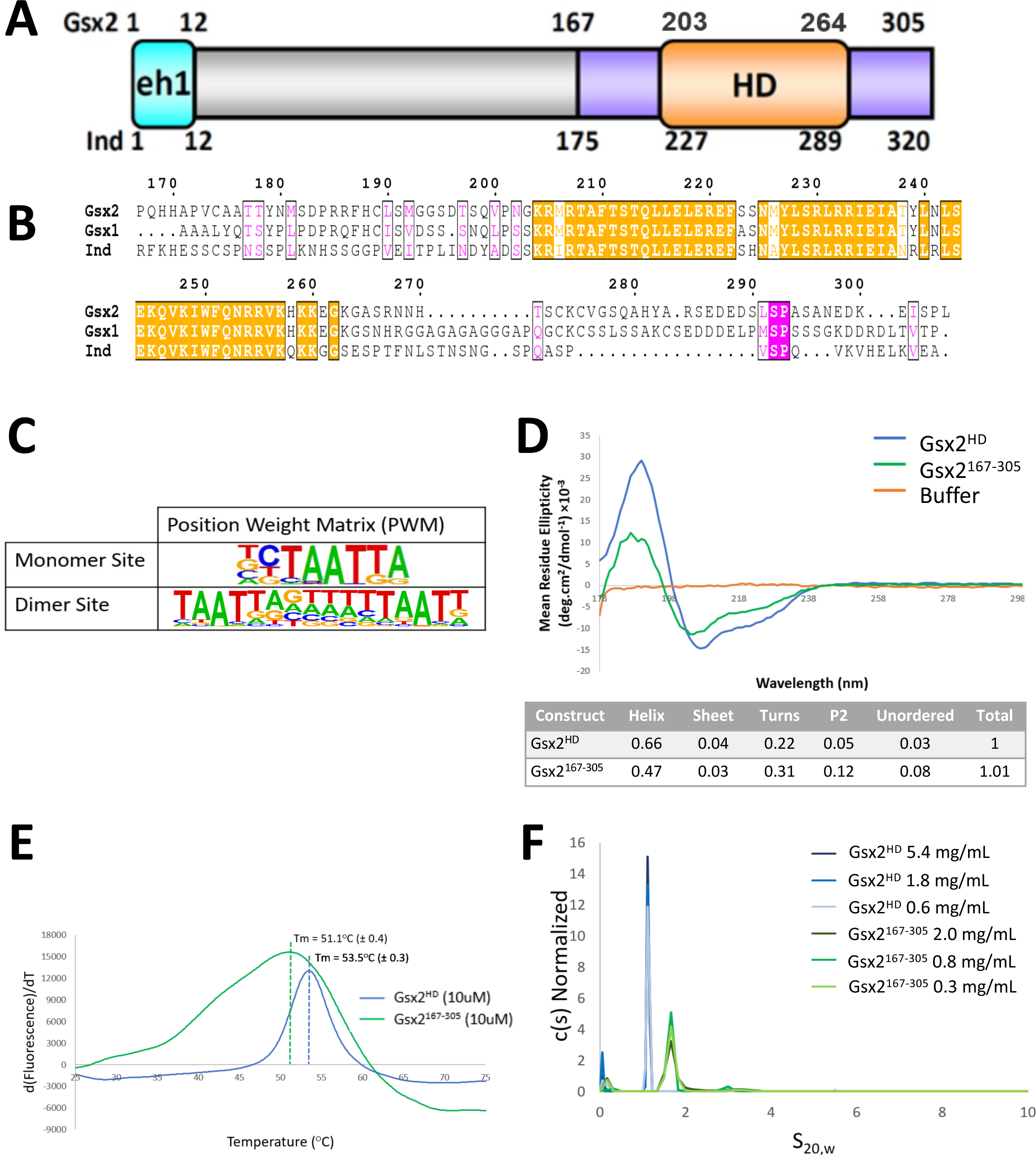
The homeodomain region of Gsx2 that cooperatively binds DNA is monomeric in solution. (A) Schematic of Gsx2 with the conserved homeodomain colored orange, flanking sequences in purple, and the eh1 repression domain in cyan. Numbers denote amino acid positions in the *M. musculus* Gsx2 and *D. melanogaster* Ind homologs. (B) Sequence alignment of the murine Gsx2^167–305^ protein with similar regions from its mouse paralog Gsx1, and its ortholog Ind from *Drosophila*. (C) The Gsx2 PWMs generated from HT-SELEX data revealed specific DNA monomer and DNA dimer sites^4,18^. (D) Far UV CD spectra depicting the difference in spectra between Gsx2^HD^, Gsx2^167–305^, and buffer. Table generated by the CDSSTR program from the DicroWeb online server estimating the secondary structure predictions of the corresponding CD data. (E) DSF assays of Gsx2^HD^ and Gsx2^167–305^ at 10 µM, highlighting a single melting peak averaging at 53.5°C and 51.1°C, respectively. (F) SV-AUC of Gsx2^HD^ and Gsx2^167–305^ at increasing concentrations yields a sedimentation coefficient distribution showing a single peak for both constructs, which corresponds to a Gsx2 monomer.

To better understand how Gsx2 regulates neural gene expression, Salomone et al. recently found that Gsx2 binds DNA as both a monomer and cooperative homodimer *in vitro* and *in vivo*^18^. Gsx2 monomers bound to a TAATTA consensus sequence, whereas dimer formation required two TAAT sequences oriented in a head-to-tail manner with a strict seven-base pair spacing between sites^18^ (Figure 1C). *Drosophila* Ind and mammalian Gsx1 were also found to have both monomer and cooperative DNA binding capabilities on similarly oriented and spaced binding sites^18,19^. Thus, cooperative DNA binding is a conserved feature of the Gsx/Ind family of proteins, suggesting that a novel mode of TF complex formation increases Gsx-specific binding to promoter and enhancer sites *in vivo*^18^. However, the molecular basis that underlies the cooperative binding of Gsx factors to DNA is not well understood.

In this study, we investigated the mechanisms used by Gsx2 to interact with DNA through a structural, biophysical, and biochemical characterization of purified Gsx protein constructs. We used analytical ultracentrifugation (AUC) and circular dichroism (CD) to show that Gsx proteins are folded monomers in solution in the absence of DNA, and isothermal titration calorimetry (ITC) to reveal that Gsx proteins bind DNA with high affinity. We used X-ray crystallography to solve the high-resolution structure of the Gsx2 HD bound to a monomer DNA site, which revealed a significant protein-dependent bend in the DNA. We incorporated the specific Gsx2 protein-DNA contacts and DNA bending characteristics from the monomer structure to build a Gsx2/Gsx2/DNA dimer model on optimally spaced DNA binding sites. This model made two key predictions: *First*, flexible A/T-rich DNA spacer sequences are preferred to allow sufficient DNA bending for the Gsx2 HD proteins to mediate direct protein-protein interactions; and *second*, specific residues on the surface of Gsx2 molecules are required to mediate cooperative DNA binding. We tested and validated both predictions using bioinformatic, biochemical, and site-directed mutagenesis approaches. Given the conservation of the key residues involved in dimer formation amongst Gsx family members, our studies identify a conserved mechanism that yields highly specific DNA binding.

## RESULTS

### Biophysical analysis reveals purified recombinant Gsx proteins are monomeric in solution

Gsx/Ind proteins have two well-defined domains; an N-terminal engrailed homology 1 (EH1) domain involved in transcriptional repression and a highly conserved C-terminal homeodomain (HD) required for DNA binding (Figure 1A-B)^18,20^. Recent biochemical studies have shown that Gsx2 can bind to DNA both as an independent monomer to individual HD sites and as a cooperative dimer on sites separated by seven base pairs (Figure 1C)^18^. To better understand the mechanisms used by Gsx2 to bind DNA, we first used bioinformatic analyses of the Gsx2 primary sequence, including AlphaFold^21^, and found that regions outside of the HD are largely predicted to be disordered (Figure S1). Thus, we focused our studies on the biophysical and structural characterization of purified recombinant proteins consisting of the Gsx2 HD (Gsx2^HD^) and a Gsx2 construct containing the HD plus short N- and C-terminal flanking regions (Gsx2^167–305^) that were shown to increase cooperativity^18^.

To assess the folding of Gsx2 in the absence of DNA, we used circular dichroism (CD), and as expected, the CD spectra for both Gsx2^HD^ and Gsx2^167–305^ showed discernable minima at 208 and 222 nm, characteristic of α-helix secondary structure content and the canonical HD fold (Figure 1D). We analyzed the Gsx2 CD data using DichroWeb^22^, which showed that both Gsx2^HD^ and Gsx2^167–305^ have high percentages of α-helix (66% and 47%, respectively). While Gsx2^HD^ has a low percentage of all other forms of secondary structure, Gsx2^167–305^ has increased percentages for disordered (8%) and polyproline II helical (P2, 12%) regions, which is expected given the predicted intrinsically disordered regions flanking the HD (Figure 1D and S1). To assess the thermostability of the Gsx2 constructs, we conducted differential scanning fluorimetry (DSF), which revealed a single melting temperature (Tm) of 54°C for Gsx2^HD^ and 51°C for Gsx2^167–305^, consistent with a thermally stable folded protein (Figure 1E). The melting curve for Gsx2^167–305^ is broader with a higher fluorescence signal than Gsx2^HD^, which is likely due to the additional ∼80 amino acid regions that flank the HD being intrinsically disordered.

Next, we used sedimentation velocity analytical ultracentrifugation (SV-AUC) to determine the oligomerization state of both Gsx2 proteins in the absence of DNA. Three samples ranging in concentration were used for both constructs; however, it should be noted that Gsx2^167–305^ concentrations were slightly lower than those used for the Gsx2^HD^ due to expression and purification difficulties but remained well above likely physiological concentrations. Figure 1F shows distributions from the sedimentation analysis. At each concentration for Gsx2^HD^, a single peak near 1 S was observed, with a frictional ratio (*f*/*f*_0_) of ∼1.4, indicating a globular protein with an estimated molar mass of 8.3 kDa – similar to the expected monomer mass of 7.9 kDa. At each concentration for Gsx2^167–305^, a single peak near 1.8 S was observed, with a *f*/*f*_0_ of ∼1.5, which is also indicative of a largely globular protein but starting to shift to a more elongated structure, again likely due to the disordered flanking regions. The estimated molar mass for Gsx2^167–305^ was 17.6 kDa, which is similar to the expected mass of 17.2 kDa. Altogether, these data support a model whereby Gsx2 proteins are folded monomers in solution.

### ITC binding studies reveal high affinity Gsx2 interactions with monomer DNA

To better understand Gsx2 interactions with DNA, we used isothermal titration calorimetry (ITC) to determine the affinity and thermodynamic parameters of both Gsx2^HD^ and Gsx2^167–305^ binding to a 15mer oligomeric DNA duplex containing a single binding site. Based on the previously determined Gsx2 monomer consensus binding site (Figure 1C) and studies of other HD protein binding sites^5,23^, we designed two 15mer oligomeric DNA duplexes that contain the sequences-TAATTA- and -TAATGG-, with -TAATTA-being the preferred site for Gsx proteins. As shown in Figure 2A and Table 1, Gsx2^HD^ tightly binds the - TAATTA- and -TAATGG-duplexes with ∼15 nM and ∼36 nM affinity, respectively, with a 1:1 stoichiometry. At 20°C, Gsx2^HD^ binding to both DNAs is enthalpically and entropically driven (Table 1). For comparison, we measured the affinity of the paralog Gsx1^HD^ on the same DNA sequences. As expected, due to its high level of sequence conservation with Gsx2 (Figure 1B), the Gsx1 HD bound similarly to the -TAATTA- and - TAATGG-duplexes with ∼18 nM and ∼42 nM affinity, respectively, and with similar enthalpic and entropic contributions to binding (Figure 2B and Table 1). However, when we determined the binding characteristics of the longer Gsx2^167–305^ protein to the two DNA duplexes, we found a decrease in its affinity for both sites compared to Gsx2^HD^. Gsx2^167–305^ has an ∼77nM affinity for the -TAATTA-site and an ∼143nM affinity for the -TAATGG-site (Figure 2C and Table 1) with two-tailed P values of 0.025 and 0.04, respectively, when compared to Gsx2^HD^. Interestingly, whereas the Gsx2^HD^ protein had nearly equal enthalpic and entropic contributions to the binding, Gsx2^167–305^ binding to both DNA sequences was nearly entirely enthalpically driven (Table 1). These findings potentially indicate that the flanking regions of Gsx2^167–305^ become structurally ordered upon DNA binding, thus incurring an entropic penalty, but do not directly contribute to DNA interactions, resulting in lower overall affinity.

**Figure 2.**
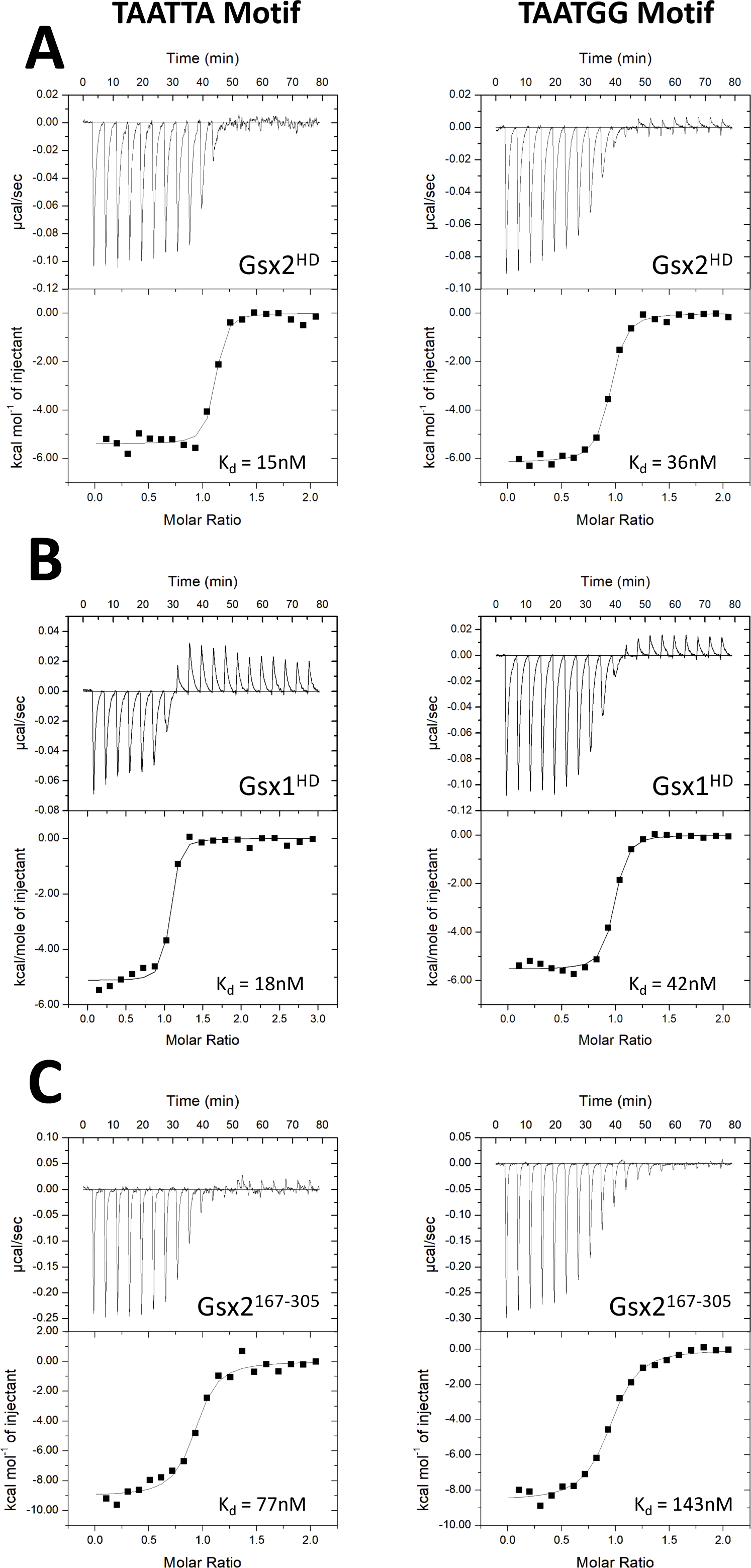
Gsx proteins bind a consensus DNA binding motif with high affinity. Isotherms from isothermal titration calorimetry (ITC) depict the binding activity of Gsx2^HD^, Gsx1^HD^, and Gsx2^167–305^ to both a -TAATTA- and -TAATGG-motif. (A) The Gsx2^HD^ shows an average 15 nM affinity for the -TAATTA-site and a 36 nM affinity for the -TAATGG-site. (B) The Gsx1^HD^ shows nearly identical affinities to Gsx2^HD^, with an average 18 nM and 42 nM affinity for the -TAATTA- and -TAATGG-sites, respectively. (C) Conversely, the Gsx2^167–305^ protein containing short N- and C-terminal flanking regions shows weaker affinity with 77 nM and 143 nM affinity for the -TAATTA- and -TAATGG-sites, respectively.

**Table 1.**
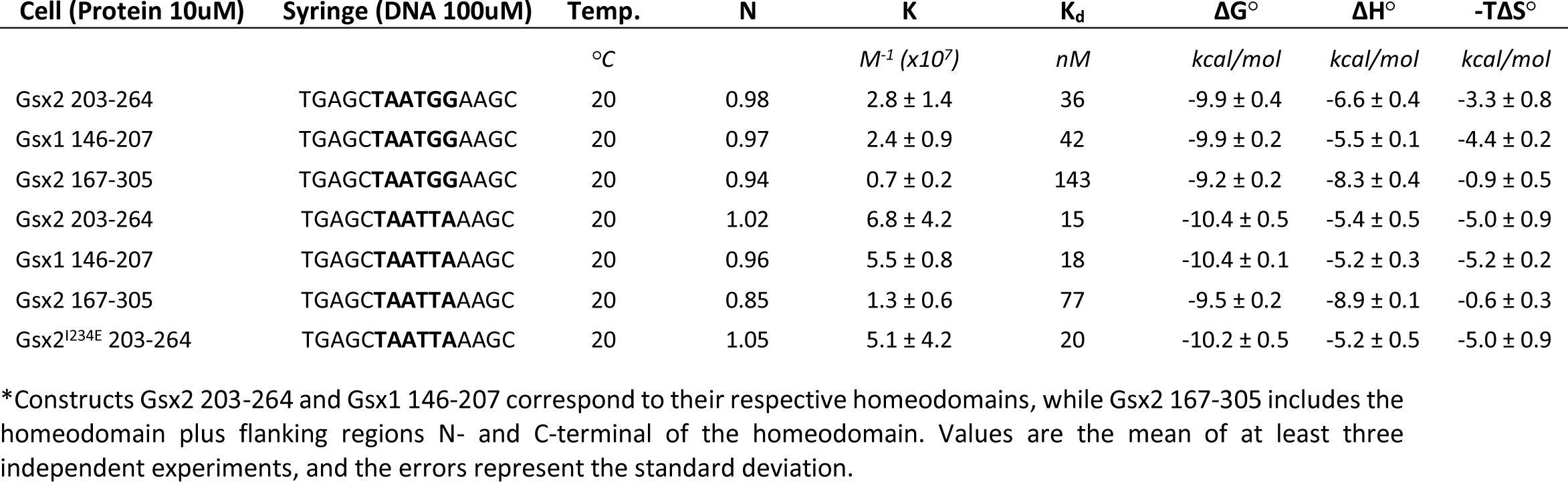
Calorimetric data of Gsx2 and Gsx1 HD constructs binding to DNA monomer sites.

### X-ray structure of Gsx2^HD^ bound to a monomer site reveals specific Gsx2-DNA interactions and DNA bending

To determine the structural basis for Gsx2 binding to DNA, we solved the X-ray structure of Gsx2^HD^ bound to a 15mer DNA containing the -TAATTA-consensus motif (Figure 3 and Table 2). The Gsx2^HD^ protein was mixed with an equal molar concentration of the 15mer duplex, containing single-stranded 5’ T and A overhangs on the sense and antisense strands, respectively. The Gsx2^HD^-DNA complex was purified by size exclusion chromatography and concentrated for crystallization screening trials. The Gsx2^HD^-DNA complex crystallized in a solution containing 30% polyethylene glycol (PEG) at pH 7.5 and crystals nominally diffracted to 2.2 Å resolution at a synchrotron X-ray source. The crystal structure of the Gsx2^HD^-DNA complex was determined using molecular replacement and refined to 2.2 Å with a R_work_/R_free_ of 22.04/26.45% (Table 2). The resulting electron density is continuous for all nucleotides, except for the two single-stranded overhangs, and is continuous for all the Gsx2^HD^ residues except for the N- and C-terminal ends (residues 203-204 and 263-264). There are two Gsx2^HD^-DNA complexes within the asymmetric unit of the crystals, forming a butterfly-like shape (Figure S2), which display a high degree of structural similarity. Alignment of all atoms (1071) of the two Gsx2^HD^-DNA complexes of the asymmetric unit results in an RMSD value of 0.265 Å (Figure S2). It should be mentioned that crystallization trials of the Gsx2^HD^ protein with DNA containing the -TAATGG-motif, as well as Gsx2^167–305^ with either DNA duplexes, were unsuccessful.

**Figure 3.**
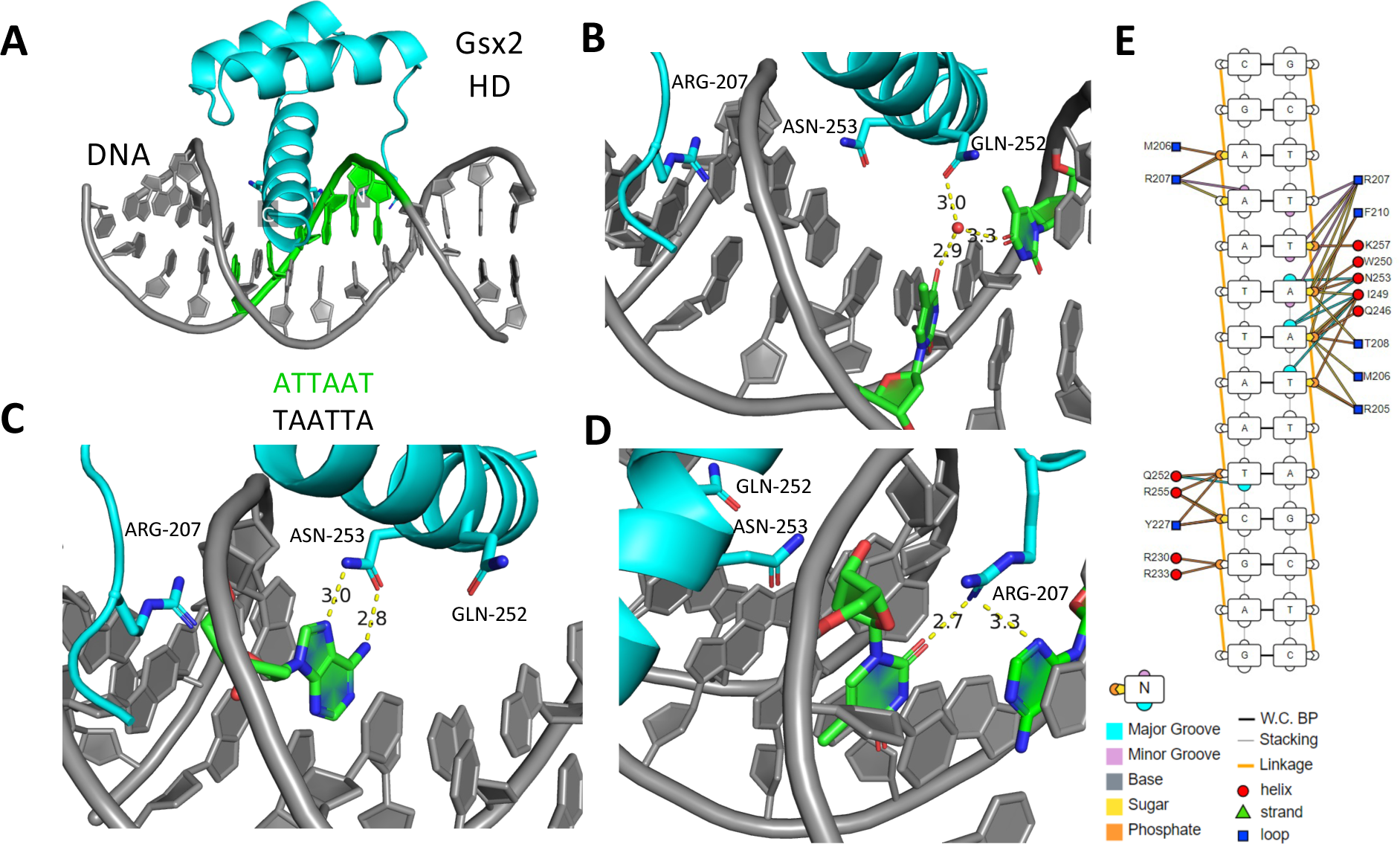
Gsx2 forms a canonical HD fold with three major contacts within the major and minor DNA grooves. X-ray crystal structure of Gsx2 203-264 (HD) bound to DNA containing the consensus motif TAATTA. (A) A single Gsx2^HD^-DNA complex shows the classic HD three helix fold, with the third helix inserted into the major groove of DNA. Gsx2 is colored cyan while DNA is colored grey, except for the - TAATTA-motif, which is colored green. N- and C-terminal ends are labeled. (B) A water molecule mediates hydrogen bond interactions between Q252 (Q50) of the HD and two thymine bases within the major groove. (C) Highly conserved N253 (N51) makes two direct hydrogen bonds with an adenine in the major groove. (D) R207 (R5) makes two direct hydrogen bonds with a thymine and an adenine in the minor groove. (E) Schematic of all specific and nonspecific protein-DNA interactions.

**Table 2.** Data collection and refinement statistics of Gsx2-DNA monomer structure.

| Data Collection Statistics |  |
| --- | --- |
| Beam line | APS LS-CAT 21-ID-G |
| Resolution (Å) | 37.43–2.20 (2.27–2.20) |
| Space group | P21 |
| Wavelength (Å) | 0.97857 |
| Unit cell a, b, c (Å) | 37.70, 37.65, 107.87 |
| Unit cell $\alpha$, $\beta$, $\gamma$ (°) | 90.00, 93.99, 90.00 |
| R<sub>merge</sub> | 0.108 (0.739) |
| I/σI | 5.1 (1.2) |
| CC<sub>1/2</sub> | 0.990 (0.405) |
| Completeness (%) | 99.7 (97.2) |
| Redundancy | 3.4 (2.7) |
| Average mosaicity (°) | 0.48 |
| Refinement Statistics |  |
| R<sub>work</sub>/R<sub>free</sub> (%) | 22.04/26.45 |
| Number of reflections | 15,200 |
| Number of atoms | 2,123 |
| Complexes/asymmetric unit | 2 |
| Wilson B/mean B value (Å<sup>2</sup>) | 44.93/54.82 |
| RMSD bond lengths (Å) | 0.013 |
| RMSD bond angles (°) | 1.514 |
| Ramachandran (favored/allowed/outliers; %) | 95.5/4.50/0.00 |

\*Highest resolution shell is shown in parentheses.

As expected, Gsx2 forms a canonical HD fold composed of three α-helices, the third of which, known as the recognition helix, fits within the major groove and makes specific and nonspecific contacts with the DNA (Figure 3A-C). Also, the Gsx2 HD has an N-terminal ARM (arginine-rich motif) that lies along the minor groove to make specific and nonspecific contacts with the DNA (Figure 3A-D). Similar to other HD structures, there are three major points of interaction between Gsx2^HD^ and DNA. First, the glutamine at the canonical 50^th^ position (Q252) of the HD forms water-mediated hydrogen bonds within the major groove to specifically interact with the thymine base at the fifth position of the -TAAT**<u>T</u>**A-motif on the sense strand and the thymine base at the sixth position of the -ATTAA**<u>T</u>**-motif on the antisense strand (Figure 3B). Second, an asparagine residue at the canonical 51^st^ position (N253) of the HD, which is conserved amongst HDs and is essential for DNA binding^2,24,25^, forms a specific bipartite hydrogen bond interaction with the adenine base at the third position of the -TA**<u>A</u>**TTA-motif (Figure 3C). Finally, the arginine residue at the canonical 5^th^ position (R207) of the HD, within the conserved N-terminal ARM of the Gsx2 HD, lies within the minor groove and makes two direct hydrogen bonds, one with the thymine base at the first position of the -**<u>T</u>**AATTA-motif on the sense strand and the other with the adenine base one position outside of the core motif -**<u>A</u>**ATTAAT-on the antisense strand (Figure 3D). Additionally, I47 (I249) is the only other residue within the major groove of the DNA that makes Van der Waals contacts with the adenine and thymine bases at the third and fourth positions, respectively, of the -TA**<u>AT</u>**TA-motif on the sense strand (Figure 3E). Numerous other nonspecific interactions between the Gsx2 HD and DNA occur along the recognition helix that are directly interacting with the phosphodiester and ribose sugar backbone of the DNA (Figure 3E). These types of interactions are common among HDs and promote the overall stability of the Gsx2^HD^-DNA complex. Collectively, these major and minor groove interactions of Gsx2^HD^ bury ∼825 Å^2^ of surface area at its interface with DNA and work synergistically to allow the HD to clamp around the sense strand of the DNA macromolecule.

To investigate how Gsx2^HD^ binding to DNA affects the structure of DNA, we used the Curves+ webserver to analyze our Gsx2^HD^-DNA X-ray structure and determine the base pair axis parameters, helical axis bending, intra- and inter-base pair parameters, backbone parameters and major/minor groove parameters (Figure 4A and Table S1)^26^. The most striking deviation is the ∼20^○^ bend observed in the DNA due to Gsx2 binding that is centered around the N-terminal ARM within the DNA minor groove (Figure 4A-B). For comparative purposes, we generated an ideal B-form of the same DNA sequence contained within our crystals using COOT^27^, which clearly highlights the degree to which the DNA is bent (Figure 4C). Additionally, we observed other alterations in the DNA parameters due to Gsx2 binding (Table S1). The most notable of these is an increase in major groove width and decrease in major groove depth that correlates with accommodation of the recognition helix into the major groove near residue N253, and a concomitant decrease in minor groove width and an increase in minor groove depth (Table S1).

**Figure 4.**
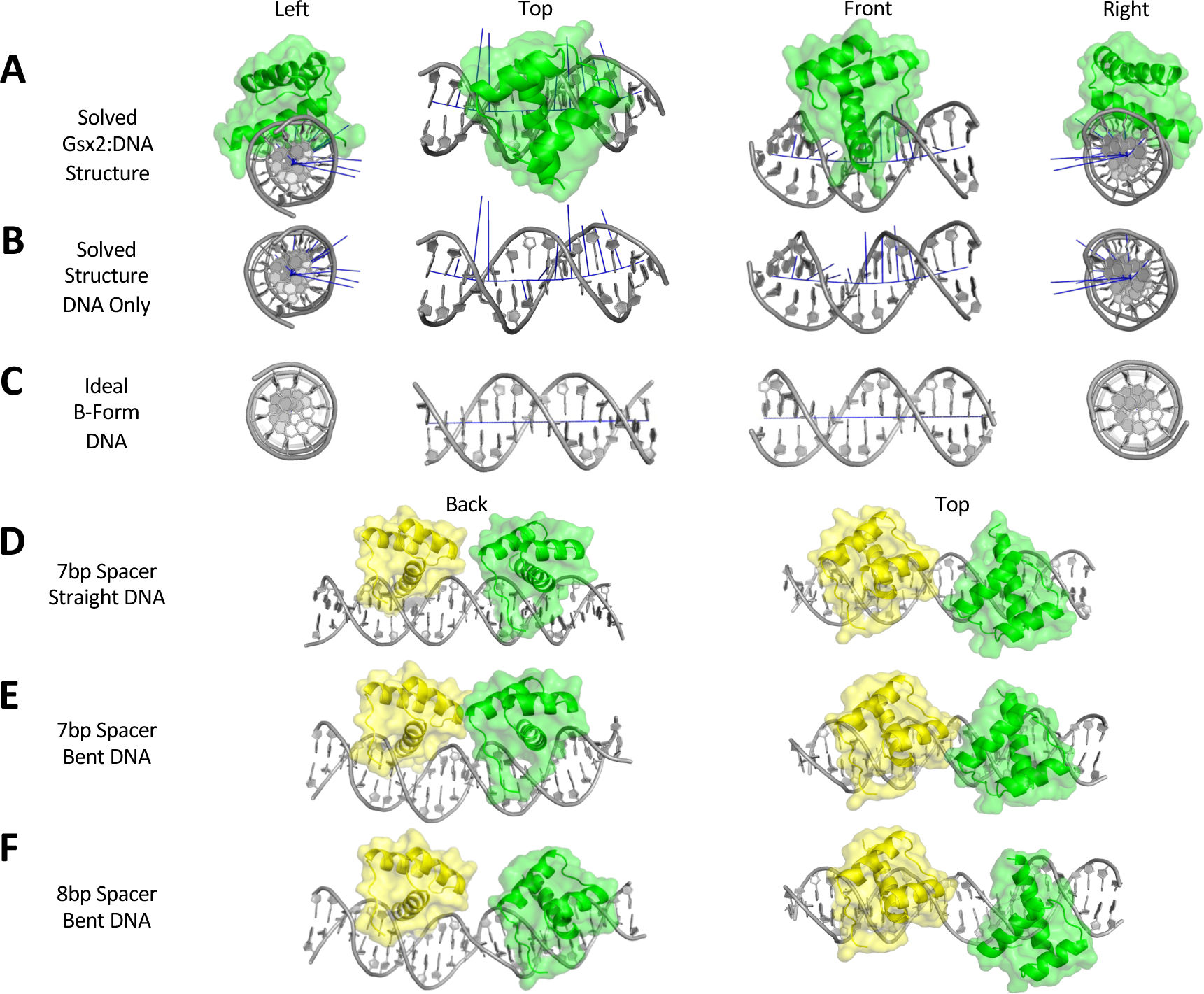
Modeling Gsx2^HD^ bound to a DNA dimer site reveals a potential protein-protein interface involved in cooperativity that is dependent on DNA bending. (A) Top, side, and bottom views of the Gsx2^HD^-DNA monomer structure, which shows significant bending of the DNA to an approximately 20° angle due to Gsx2^HD^ interactions. Gsx2^HD^ is shown in green, and DNA is shown in light grey. Blue lines parallel to the DNA represent the helical axis, while blue lines perpendicular to the DNA represent the degree and directionality of DNA bending. (B) Isolated view of the DNA from the structure without the Gsx2 protein, highlighting DNA bending. (C) Comparison views of an ideal B-form DNA duplex with an identical sequence to the DNA used in the Gsx2^HD^-DNA structure. (D) Top and back view of a Gsx2^HD^-DNA dimer model with optimal dimer sequence, 7bp spacer length, and orientation, but lacking any DNA bending. One Gsx2^HD^ protein is in yellow, the second Gsx2^HD^ protein is in green, and the DNA is light grey. No protein-protein contacts are observed in this model. (E) Comparison views of a Gsx2^HD^-DNA dimer model based upon a dimer sequence with 7bp spacer length and 20° bend observed in monomer structure. Potential protein-protein interactions are observed between both Gsx2^HD^ proteins. (F) Comparison views of a Gsx2^HD^-DNA dimer model with a dimer sequence with an 8bp spacer length and DNA bending shows a loss of direct contact between the two Gsx2^HD^ proteins.

### Modeling Gsx2 binding to a DNA dimer site reveals DNA bending and a novel protein-protein interface that are required to facilitate cooperative binding

To gain insight into the Gsx2 protein-protein interactions that underlie cooperative binding to a DNA dimer site, we used our structure of the Gsx2^HD^ bound to a DNA monomer site to create a dimer model of two Gsx2^HD^ proteins bound to a DNA dimer site with the correct seven base pair spacing and head-to-tail orientation between sites (Figure 1C). To do so, we first used the modeling software PyMol (The PyMOL Molecular Graphics System, Version 2.0 Schrödinger, LLC) to trim nucleotide base pairs from both ends of the monomer structure. The base pairs removed were specifically selected to have the smallest effect on DNA bending and maintain the DNA curvature observed within the Gsx2^HD^-DNA monomer structure (Figure 4A-B). The DNA ends of the two monomer structures were subsequently joined to maintain correct bond lengths and angles within the phosphodiester backbone of DNA, and the degree of DNA bending predicted by this Gsx2^HD^-DNA dimer model was validated with Curves+^26^. Strikingly, this model predicts that the two Gsx2^HD^ proteins directly contact each other without clashes (Figure 4E).

To assess whether DNA bending and spacer length between sites were critical parameters for the formation of the predicted protein-protein interactions between Gsx2^HD^ molecules, we built two additional Gsx2^HD^/DNA dimer models: First, we modeled the Gsx2^HD^ on ideal B form DNA (i.e., no DNA bending) and found that the two Gsx2^HD^ proteins would not interact in the absence of DNA bending (Figure 4D). Second, we modeled the Gsx2^HD^ on a DNA dimer site with an eight base pair spacing, which has previously been shown to abrogate cooperative binding^18^, and found that the two Gsx2^HD^ proteins would not interact due to both the increased distance/separation and the altered rotation/periodicity of the DNA causing a misalignment of the monomers (Figure 4F). Taken together, these modeling studies suggest that Gsx2^HD^ binding to the individual TAAT sequences induce DNA bending and thereby promotes the formation of a protein-protein interaction interface between the two Gsx2^HD^ molecules that stabilizes cooperative homodimer binding.

### Testing the role of DNA spacer flexibility in cooperative Gsx2 DNA binding

Given that our Gsx2^HD^-DNA dimer model suggested DNA bending was important for cooperative binding, we first analyzed whether the identity and flexibility of the spacer sequence between the two Gsx2 binding sites impacts cooperative DNA binding. Since prior studies have shown that adjacent DNA sequences not directly contacted by a TF can impact the flexibility of DNA^28^, we reasoned that Gsx2 may prefer flexible DNA sequences between the two sites to better enable DNA bending and dimer formation.

Consistent with this idea, analysis of the published Gsx2 dimer binding motif consensus sequence determined by HT-SELEX for the spacer region between the two sites is enriched for A/T base pairs^4,18^ (Figure 5A), which are known to be more flexible than G/C sequences^29^. To assess how spacer identity impacts cooperative Gsx2 dimer formation, we compared the rate of dimer site enrichment for A/T-rich versus G/C-rich spacer sequences using the HT-SELEX data. Because the Gsx2^HD^ protein directly contacts the first two nucleotides of the spacer sequence in the -TAAT**<u>NN</u>**NNNNNTAAT-dimer site, we excluded these nucleotides from our analysis and focused on the remaining five spacer nucleotides (boxed in Figure 5A). We next determined the percentage of sequences containing dimer sites with A/T-rich spacers versus dimer sites with G/C-rich spacers as a function of HT-SELEX cycle. As expected, the number of dimer sequences with either A/T or G/C spacers is similar and very low in the initial library (HT-SELEX cycle 0) and after a single round of binding (HT-SELEX 1) (Figure 5A, bottom). Strikingly, however, the number of dimer sequences with A/T-rich spacers dramatically increases compared to dimer sequences with G/C-rich spacers after cycles 2 through 4. These data suggest that Gsx2 binding preferentially enriches for dimer probes with A/T-rich spacer sequences over G/C-rich spacer sequences.

**Figure 5.**
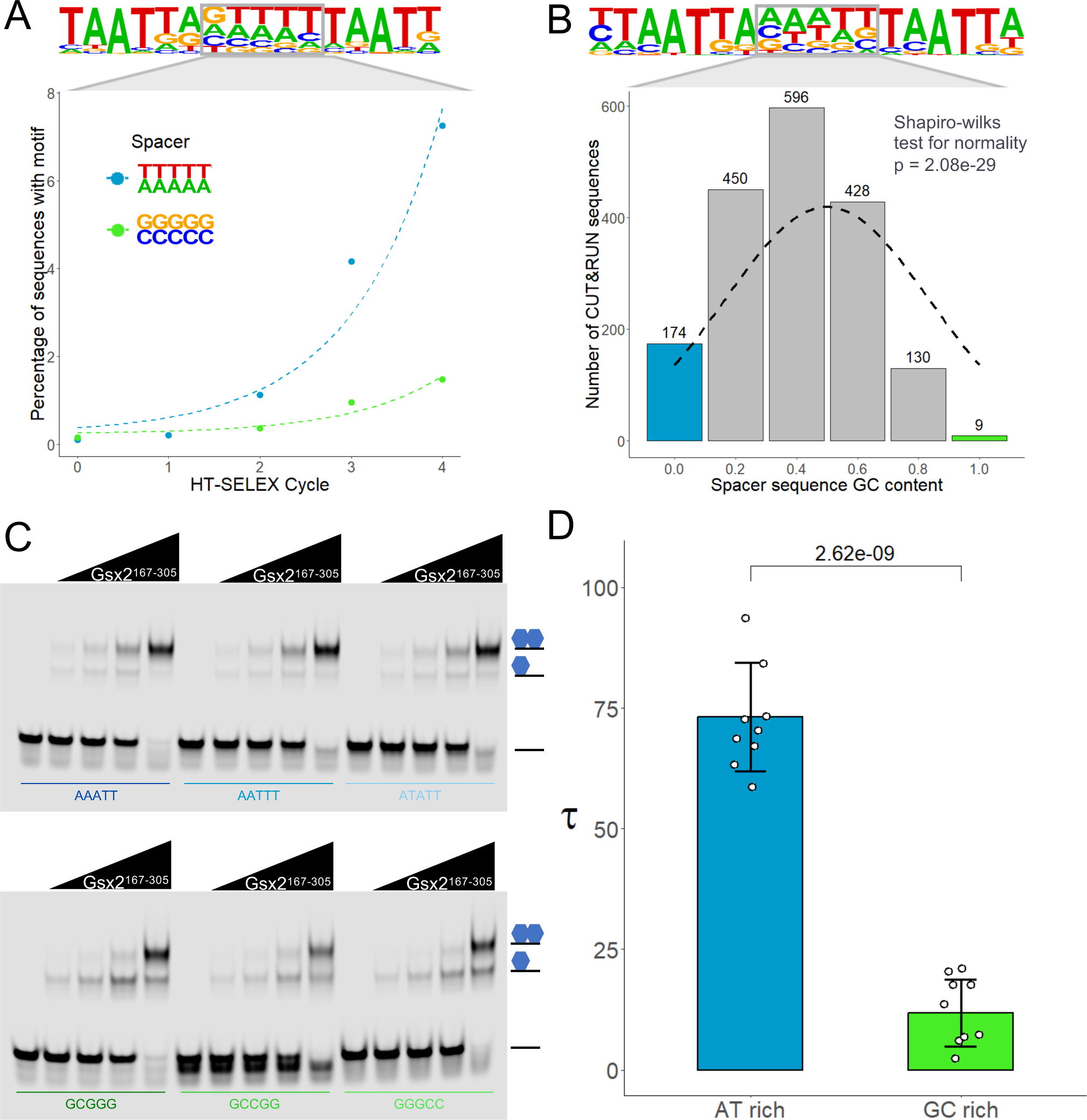
Gsx2 preferentially selects DNA dimer sequences with flexible A/T-rich spacers. (A) Analysis of *in vitro* HT-SELEX data^4^ revealed significantly faster enrichment for dimer sequences with A/T-rich spacers compared to G/C-rich spacers. Note, the PWM generated from analyzing the sequences after the 4^th^ round of selection at top reveals a boxed spacer sequence of 5 nucleotides with strong overall A/T preference. Graph below depicts the percentage of sequences encoding A/T versus G/C rich spacers in the original library (cycle 0) and after each round of selection (cycles 1 through 4). (B) Analysis of *in vivo* CUT&RUN data showing that Gsx2 has a bias for DNA dimer sequences with A/T-rich spacers. At top is the Gsx2 dimer PWM from the top quartile of called peaks and the dashed line in the graph below indicates a perfect Gaussian curve if no bias was detected. (C) EMSAs of Gsx2^167–305^ binding to DNA dimer duplexes with various A/T-rich or G/C-rich spacers show increased cooperative binding for all A/T-rich sequences compared to G/C-rich sequences. Note, each EMSA probe was tested in 5 lanes containing the following concentrations of purified Gsx2^167–305^ protein (0, 50, 100, 200, and 400 nM) (D) The cooperativity of the EMSAs was quantified using the Tau factor, which reveals an approximately 6-fold increase for A/T-rich sequences versus G/C-rich sequences, consistent with our bioinformatic data highlighting Gsx2’s preference for more flexible A/T-rich spacer sequences. Each dot represents the Tau value from an individual binding reaction of either an A/T rich or G/C rich spacer probe (n = 12 for each group). Error bars denote standard deviation. Tau factors were compared with a two-sided unpaired student t-test.

To determine if Gsx2 also preferentially binds dimer sites with A/T-rich spacer sequences *in vivo*, we analyzed available CUT&RUN genomic binding data for Gsx2 from the mouse forebrain^18^. CUT&RUN assays provide a sensitive method to detect *in vivo* DNA binding, and notably, the *in vivo* CUT&RUN PWM for mouse Gsx2 dimer sites closely matches the *in vitro* HT-SELEX PWM for human GSX2 with the spacer sequence similarly enriched for A/T sequences (compare Figure 5A and 5B). To quantify the *in vivo* spacer sequence preference of Gsx2, we determined the A/T versus G/C content across all five spacer nucleotides using the experimentally identified genomic dimer sites (∼1,800 sites) and found that the sequence distribution was significantly skewed towards A/T-rich sequences compared to the expected normal distribution (Figure 5B). Thus, the A/T-rich spacer sequence bias between Gsx2 binding sites was observed in both *in vitro* and *in vivo* DNA binding assays.

Next, we experimentally tested if Gsx2 cooperativity preferentially occurs on dimer sites with A/T-rich spacers using quantitative EMSAs to measure the cooperativity of Gsx2 binding to six different DNA dimer probes: three of which contained different A/T spacers and three of which contained different G/C spacers (Figure 5C). All of the remaining nucleotides were kept the same between probes and we measured cooperativity from the EMSAs by calculating the Tau factor as previously described^30^ (see Methods). As shown in Figure 5C and D, while Gsx2^167–305^ formed cooperative dimer complexes on both A/T and G/C spacer probes, Gsx2 much more readily formed dimer complexes on the A/T spacer probes (Tau = 72.5) compared to the G/C spacer probes (Tau = 12.6). These data suggest that the composition of the spacer sequence affects cooperativity independent of direct DNA-protein interactions with Gsx2. Moreover, it is important to note that Gsx2 similarly depleted each free probe in these EMSAs (Figure 5C; Figure S3B), suggesting that the different spacer sequences did not significantly affect the affinity of Gsx2 for the individual binding sites. To further analyze these spacer sequences, we used DNA shape prediction software with each of our probes (Figure S3C-F). Previously, it has been shown that the large dipole present in G-C pairs produces electrostatic interactions that are not conducive to base pair stacking, which decreases the compressibility of the major and minor grooves, thereby reducing the flexibility of the DNA^29^. Consistent with A/T-rich sequences exhibiting increased DNA flexibility, the minor groove width of the A/T-rich probes was substantially smaller than the G/C probes (Figure S3C), and the propellor twists and roll of the A/T-rich probes had greater magnitudes when compared to the G/C probes (Figure S3D-E). Taken together, the Gsx2 HT-SELEX data, genomic binding data, and quantitative EMSAs show that the ability of Gsx2 to bind DNA dimer sites cooperatively is significantly influenced by the composition of the spacer DNA sequences between the two sites, i.e., cooperative Gsx2 binding to dimeric DNA is greatly enhanced by A/T rich spacer sequences that bend more readily compared to G/C rich sequences.

### Identification of a Gsx2-Gsx2 interface required for cooperative DNA binding

In addition to DNA bending, the Gsx2^HD^-DNA dimer model also identified key residues within the Gsx2^HD^-Gsx2^HD^ interface predicted to be required for cooperative DNA binding (Figure 4E). Using this model, we identified four residues likely to contribute to cooperativity by mediating protein-protein interactions between the two Gsx2^HD^ molecules (Figure 6A). Gsx2 chain A residues L231 and I234 (colored yellow in Figure 6A), which reside at the beginning of the second α-helix in the Gsx2^HD^ structure, and Gsx2 chain B residues S212 and L216 (colored green in Figure 6A), which reside at the beginning of the first α-helix in the Gsx2^HD^ structure, compose the modeled Gsx2(A)-Gsx2(B) interface. Interestingly, all Gsx family members, including Ind, contain these four residues, whereas closely related HD TFs lack many of these residues (Figure 6B). We designed glutamate mutations at these four sites (S212E, L216E, L231E, and I234E) to compare the cooperative binding of WT and mutant Gsx2^HD^ constructs to DNA. Equimolar concentrations of each protein were tested in EMSAs with DNA probes containing either the optimal 7bp spacer dimer site (7bpS) as defined by HOMER analysis of available HT-SELEX data^4^ or this same sequence with an additional base pair to generate a sub-optimal 8bp spacer (8bpS) site. As shown in Figures 6C and S4, comparative EMSA analysis of both Gsx2^HD^ wild-type (denoted WT^HD^ in Figure 6C) and mutants revealed that all four mutations reduced the ability of Gsx2^HD^ to cooperatively dimerize on DNA with I234E having the most dramatic impact. Importantly, each mutation specifically affected cooperativity without generally affecting DNA binding as judged by the relative amounts of free DNA probe remaining in each EMSA lane when comparing WT^HD^ to mutants (Figures 6C and S4). Given that I234E (I234E^HD^) had the most pronounced effect on cooperativity, we performed EMSAs in triplicate with I234E^HD^ and WT^HD^ for Tau analysis of cooperativity. As expected, Gsx2 WT^HD^ is cooperative on the 7bpS probe, but not on the 8bpS probe, with Tau factors of ∼38 and ∼2, respectively (Figure 6C; Figure S5A-C). In stark contrast, the I234E^HD^ mutant had Tau factors of ∼1.6 for both the 7bpS and 8bpS probes, which is ∼24-fold less cooperative than WT^HD^ (Figure 6C). Moreover, we used ITC to ensure that the I234E mutation did not generally affect Gsx2^HD^-DNA interactions and found virtually identical monomer DNA binding affinities between the I234E^HD^ and WT^HD^ proteins (Table 1 and Figure S6). Finally, due to the higher cooperative binding seen previously with the longer Gsx2^167–305^ protein compared to Gsx2^HD^, we performed similar comparative EMSAs with WT and I234E Gsx2^167–305^ proteins and found that the I234E mutation similarly compromised cooperative binding to the 7bpS probe (Figure 6D). Taken together, these studies reveal that Gsx2 uses a novel protein-protein interface to mediate cooperative binding to DNA containing a dimer site, and altering residue I234 disrupts this binding interface.

**Figure 6.**
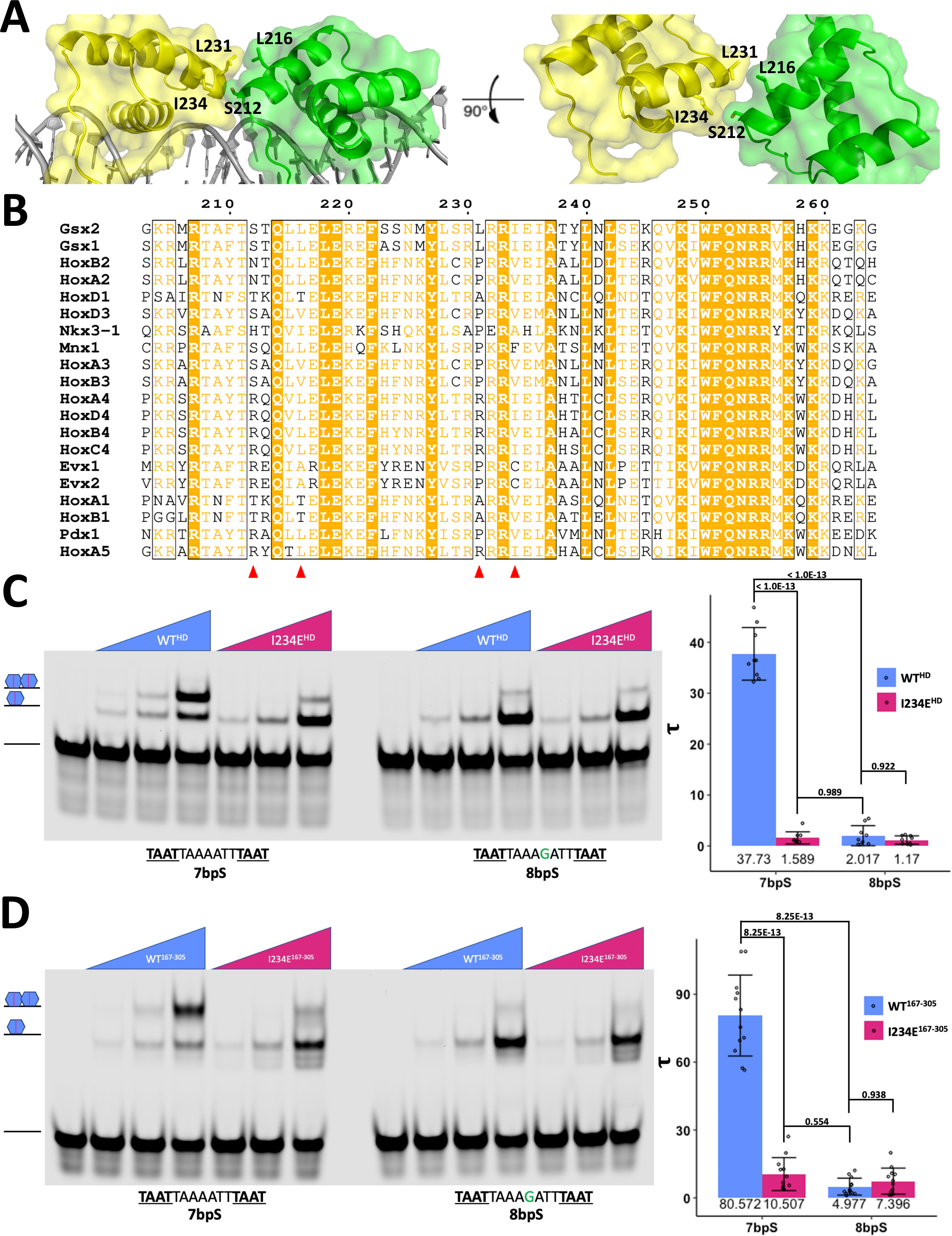
Gsx2^HD^ dimer model reveals a novel binding interface with residue conservation amongst the Gsx/Ind family. (A) Expanded front and top view of the protein-protein interface observed in our 7bp spacer length dimer model. Four residues are primarily localized at the interface: S212, L216, L231, and I234. The two Gsx2^HD^ monomers are colored yellow and green, and the interface residues are shown in stick representation. (B) Sequence alignment of the HDs Gsx2^HD^ and Gsx1^HD^, which have been shown to bind DNA cooperatively, with close relatives, none of which are expected to bind DNA cooperatively. Red triangles denote residues S212, L216, L231, and I234. (C) A representative EMSA and corresponding Tau factor calculations of WT^HD^ and I234E^HD^ binding to the 7bpS (7bp spacer, cooperative) and 8bpS (8bp spacer, non-cooperative) DNA dimer probes, demonstrating that the I234E^HD^ protein shows significantly reduced cooperative binding on the 7bpS probe compared to the WT^HD^ protein. Note, each EMSA probe was tested in 4 lanes containing the following concentrations of purified Gsx2^HD^ protein (0, 25, 100, and 400nM). Each dot represents the Tau value from for either the 7bp or 8bp spacer probe at each of the different protein concentrations. The mean Tau value for each probe:protein combination is noted and error bars denote standard deviation of the mean. Significance was determined by two-way ANOVA with Tukey’s multiple comparisons test. (D) Replicate EMSA from A but instead showing WT^167–305^ and I234E^167–305^ binding to the 7bpS and 8bpS DNA dimer probes. Again, only WT^167–305^ on the 7bpS probe is strongly cooperative, demonstrating that even in the presence of the flanking regions around the HD the I234E mutation is sufficient to greatly diminish Gsx2 cooperative DNA binding. Note, each EMSA probe was tested in 4 lanes containing the following concentrations of purified Gsx2^167–305^ protein (0, 25, 100, and 400 nM). Each dot represents the Tau value from for either the 7bp or 8bp spacer probe at each of the different protein concentrations. Numbers beneath graph bars denote the mean Tau value for each probe:protein combination. Error bars denote standard deviation of the mean. Significance was determined by two-way ANOVA with Tukey’s multiple comparisons test.

## DISCUSSION

Gsx2 and its paralog Gsx1 are part of the Hox-like class of the HD superfamily, which contains ∼40 members^1^. The vast majority of Hox-like HDs exhibit preferential affinity for highly similar A/T-rich DNA motifs *in vitro*, as conferred by their respective DNA-binding HDs^5,6,31^. An outstanding question in the field is to identify the mechanisms and interactions that promote the required *in vivo* target gene specificity to prevent the mistargeting of HD TFs. One compelling mechanism is the ability of some HD TFs to dimerize on DNA^24,25^. Previously, it was shown that Gsx factors can bind both monomer and cooperative dimer DNA sites *in vitro* and *in vivo*, thereby increasing their specificity^18^. More recently, Cain *et al.* used a bioinformatic approach to analyze HT-SELEX data to identify members of different HD subclasses (e.g., paired-like, NK-like, etc.) that also cooperatively bind DNA dimer sites with specific base pair spacers between the sites^19^.

Here, we used biochemical, biophysical, and structural approaches to better understand the mechanistic basis for Gsx2 binding to monomer DNA sites followed by molecular modeling and quantitative DNA binding assays to define the mechanisms underlying cooperative binding to dimer DNA sites. First, we used AUC studies to show that Gsx2 exists as a monomer in solution, even at concentrations well above likely physiological levels, and regardless of the presence of the flanking regions outside of its HD (Figure 1F); second, our ITC binding studies showed that Gsx2 interacts with a DNA monomer with low to mid nanomolar affinity (Figure 2); and third, we used X-ray crystallography to solve the structure of the Gsx2^HD^ bound to a DNA monomer site (TAATTA) (Figure 3). While we were unable to solve the X-ray structure of the cooperative Gsx2 dimer bound to DNA, we built and validated molecular models of the Gsx2 dimer on DNA by combining our monomeric structure with previous data defining the sequence, orientation, and spacer length requirements for DNA dimer sites^18,19^ (Figure 4). Importantly, the resulting dimer models revealed that the ∼20^○^ bend in the DNA observed in the Gsx2^HD^-DNA monomer structure is critical for creating a binding interface between the two appropriately spaced Gsx2^HD^ molecules. Moreover, the dimer model suggests that Gsx2^HD^ binding to individual monomer sites induces bending in the DNA, thereby promoting the formation of a protein-protein interface between the two Gsx2^HD^ molecules that stabilizes cooperative homodimer binding.

In support of the role of Gsx2-mediated DNA bending in cooperative dimer formation, we found that Gsx2 preferentially binds DNA dimer sites containing flexible A/T-rich 7bp spacer sequences in both *in vitro* HT-SELEX and *in vivo* CUT&RUN data (Figure 5). We experimentally validated these bioinformatic data by performing quantitative EMSAs, which showed that Gsx2 cooperativity preferentially occurs on DNA probes with A/T-rich spacer sequences over G/C-rich spacer sequences (Figure 5C-D). We also used the dimer model to identify a protein-protein interface between two Gsx2 molecules that mediates cooperative binding to dimeric DNA (Figure 6A). We subsequently used site-directed mutagenesis and quantitative EMSAs to show that mutations in all four predicted residues compromised cooperativity (Figure 6 and S4) with the I234E variant having the largest impact on cooperativity in both the short HD only protein (I234E^HD^) and the longer Gsx2^167–305^ proteins (I234^E167–305^) (Figure 6C-D). Importantly, we showed by ITC that I234E^HD^ bound monomeric DNA with virtually identical affinity as the WT^HD^ protein (Table 1 and Figure S6), suggesting that residue I234 plays an energetically pivotal role in forming the cooperative Gsx2 dimer interface on DNA, but does not generally affect interactions with DNA.

From an evolutionary standpoint, the four residues in Gsx2 that affect cooperative binding are conserved in the mammalian paralog Gsx1 and the *Drosophila* ortholog Ind (Figure 6B), which also form cooperative dimers on DNA sites with 7bp spacers^18^. This suggests that the Gsx/Ind HD interface involved in cooperative binding on DNA dimer sites is conserved from flies to humans. When we aligned Gsx2 with closely related Hox-like HD TFs based on primary sequence, some, but not all, of the interfacial residues are conserved, and none of these related HDs have all four residues conserved (Figure 6B). Interestingly, based on previous work that analyzed HT-SELEX sequencing data, none of these related HD TFs are expected to form cooperative homodimers on DNA^19^.

Previous structural studies of other HDs that form cooperative dimers on DNA have revealed that protein-protein interactions play a major role in the molecular basis of cooperativity, e.g. the Paired homodimer and Al/Cll (aristaless/clawless), Ubx/Exd (ultrabithorax/extradenticle), AbdB/Exd (abdominal B/extradenticle), and Scr/Exd (sex combs reduced/extradenticle) heterodimer structures from *Drosophila*^30,32–35^, the MATa1/MATα2 heterodimer structure from yeast^36^, and the HOXB1/PBX1 and MEIS1/DLX3 heterodimer structures from human^37^. Strikingly, however, none of these proteins cooperatively bind DNA sites with a 7bp spacer and the Gsx2/Gsx2 interface predicted from our dimer model, which includes residues at the beginning of the second α-helix of one Gsx2 HD and residues at the beginning of the first α-helix in the second Gsx2 HD, has not been observed to mediate cooperativity in any of the HD dimer structures mentioned above. These findings suggest that we have identified a novel HD-HD interface involved in cooperative binding to DNA containing a dimer site.

While our dimer model does not directly address how the flanking regions of Gsx2^167–305^ increase cooperativity compared to the isolated Gsx2^HD^ ^18^, it does reveal that the distances between the two N-or C-termini of DNA bound Gsx2 molecules are separated by ∼35 Å. Given the ∼40 residues flanking the HD on both ends, this would likely allow for additional protein-protein and/or protein-DNA interactions to increase cooperativity. However, the residues and motifs contained within the flanking regions that contribute to cooperative binding have not been finely mapped. Certainly, additional future studies are warranted, including a high-resolution structure of Gsx2 constructs containing the flanking regions bound to dimer DNA, which will be important to provide a complete structural understanding of cooperative binding by Gsx2.

## Data Availability

The structure has been deposited into the Protein Data Bank (PDB) with accession number 8EML.

## Acknowledgments

We thank Kenny Campbell and his lab for their constructive criticism and the beamline staff at LS-CAT for their technical assistance.

## Contributions

This scientific study was conceived and planned by J.W., E.G., R.K., and B.G. All construct synthesis and bacterial protein purifications were carried out by J.W. and Z.Y. All CD experiments were performed by J.W. with assistance from A.E.Y. DSF assays were performed by J.W., Z.Y., and E.G. AUC experiments were performed and analyzed by A.E.Y., E.S., and A.B.H. All ITC experiments were performed by J.W. and Z.Y. All crystallography and X-ray structural studies were conducted by J.W. and analyzed by both J.W. and R.K. Modeling of the Gsx2-DNA dimers and identification of the Gsx2-Gsx2 binding interface was performed by J.W. Bioinformatic analyses of HT-SELEX and CUT&RUN data as well as DNA shape analyses were conducted by B.C. Site-directed mutagenesis was carried out by J.W. and Z.Y. EMSAs and Tau factor calculations were performed by E.F. and B.C. The manuscript was written by J.W., B.G. and R.K. and edited by all authors.

## Materials & Methods

### Cloning, Expression, Purification

All Gsx2 constructs were subcloned from a cDNA containing the mouse *Gsx2* gene into a modified version of the pET-14b vector (Novagen) that includes an N-terminal His-tag followed by a PreScission Protease cleavage site in place of the original thrombin cleavage site. The expression vector was transformed into C41(DE3) (Sigma-Aldrich) *E. coli,* and bacteria were grown in autoinduction media^38^ at 37^◦^C for 3 hours, cooled to 20^◦^C, and then grown overnight. The cultures were harvested by centrifugation, and cell pellets were resuspended in binding buffer (1XBB; 20mM Tris 8, 500mM NaCl, 5mM Imidazole) and frozen. Frozen pellets were thawed, lysed by sonication, cleared by centrifugation, and incubated with Ni-NTA beads. Beads were then loaded into a gravity column and washed with 1XBB with 0.1% Triton and 1xBB with 0.1% NP40. Protein was eluted using 1XBB with 0.1% NP40 and 0.5M Imidazole. The eluted protein was dialyzed, and the His-tag was cleaved with PreScission Protease (Cytiva) per the manufacturer’s protocol. Gsx2 protein was further purified via cation exchange and size exclusion chromatography. Purity was assessed by SDS-PAGE gel electrophoresis followed by Coomassie staining. Finally, the Gsx2 protein was concentrated to ∼20-40 mg/ml in a buffer containing 20mM MES pH 6, 150mM NaCl, 1% ethylene glycol, and 0.1mM TCEP.

### Circular Dichroism (CD)

CD experiments were performed on an Aviv Circular Dichroism Spectrophotometer 215 using a 0.5mm quartz cuvette (Hellma Analytics). The cuvette was not removed during a series of scans taken from 300nm to 190nm in 1nm increments. Proteins were dialyzed into a buffer containing 5mM sodium phosphate pH 6.5, 150mM NaF, and then diluted to the desired concentration of ∼0.30 mg/ml in the same buffer. The resulting data were analyzed via the Dichroweb server using the CDSSTR program with reference set 5^22^. Data are plotted as mean residue ellipticity, [θ], in units of degrees cm^2^ dmol^−1^ residue^−1^.

### Analytical Ultracentrifugation (AUC)

Experiments were performed using a Beckman Coulter XL-I analytical ultracentrifuge with an An-60 Ti rotor. Data were collected using interference optics. Experiments were run at a speed of 48,000 rpm overnight until no further sedimentation was observed (approximately 20 hours). Proteins were dialyzed into a buffer containing 50 mM sodium phosphate pH 6.5, 150 mM NaCl, and 0.1 mM DTT and diluted to a desired concentration in the same buffer prior to loading the samples. The homeodomain construct was run at concentrations of 0.6, 1.8, or 5.4 mg/ml using meniscus-matching two-sector epon-charcoal 1.2-cm centerpieces (Spin Analytical) with sapphire windows. The extended construct containing the flanking sequences was similarly run at concentrations of 0.3, 0.8 and 2.0 mg/ml. Samples were equilibrated at 20 °C for at least 1 hour before beginning experiments. SEDFIT’s^39^ continuous c(s) distribution model was used to analyze data. SEDNTERP^40^ was used to estimate the partial specific volume, buffer density, and buffer viscosity. The S_20,w_ value is the sedimentation coefficient of a single species under standard conditions (20 °C in water). The S_20,w_ value, frictional ratio (f/f0), and resulting calculated molecular weights were reported by SEDFIT’s c(s) distribution analysis^39^.

### Isothermal Titration Calorimetry (ITC)

ITC experiments were performed using a Microcal VP-ITC microcalorimeter. For all experiments, the DNA duplexes were placed in the syringe at ∼100uM, and all Gsx2 proteins were placed in the cell at ∼10uM. Titrations consisted of an initial 1uL injection followed by nineteen 14ul injections. All experiments were performed in a buffer containing 50mM sodium phosphate pH 6.5 and 150mM NaCl at 20^◦^C. All samples were dialyzed overnight to ensure buffer match. Final raw data were analyzed using ORIGIN and fit to a one-site binding model.

### Crystallization

Gsx2-DNA complexes were formed prior to crystallization by mixing purified protein in a 1:1.2 (protein:DNA) ratio with a final complex concentration of ∼15mg/ml (∼900uM). The DNA for crystallization was a 15-mer duplex with single-strand overhangs, containing the sequence sense strand 5’ – TGAGC<u>TAATTA</u>AAGC – 3’ and the antisense strand 5’ – AGCTT<u>TAATTA</u>GCTC – 3’. Crystallization conditions were initially screened using the BCS screen from Hampton. The crystallization condition that gave rise to crystals was a 1:1 ratio mix of protein-DNA solution and well solution (0.1M MgCl_2_, 0.1M RbCl, 0.1M HEPES pH 7.5, 30% PEG Smear Broad). The final crystallization condition was equilibrated over 500ul of well solution and grown at 4^◦^C. Subsequent crystals diffracted to 2.2 Å and belong to the monoclinic space group P2_1_ with cell dimensions 37.70, 37.65, 107.87 Å. The asymmetric unit of the crystal contained two Gsx2-DNA complexes.

### Structure Determination, Model Building, and Refinement

Phaser^41^ was used for molecular replacement with the complex of Pdx1 and DNA (2H1K) as a search model^42^. Two Gsx2-DNA complexes were observed within the asymmetric unit of the crystal. COOT^27^ was used for manual model building within the observed electron density. Phenix^43^ was then used for general refinement and the selection of TLS parameters for additional model refinement. Finally, the model was validated with MolProbity^44^. The final model was refined to a R_work_ = 22% and a R_free_ = 26% with good overall geometry. PyMOL (The PyMOL Molecular Graphics System, Version 2.5.2, Schrödinger, LLC.) was used to create all figures of the structure.

### Modeling the Gsx2 Dimer

Dimer models were generated using PyMol (The PyMOL Molecular Graphics System, Version 2.5.2, Schrödinger, LLC.). The X-ray crystal structure of Gsx2 bound to a DNA monomer site was duplicated with minor deletions of the DNA end nucleotides to account for the precise seven base-pair and eight base-pair spacers. Deletions were carefully selected to ensure the dimer model DNA maintained the approximately 20^◦^ bend, as seen in the monomer structure. No other alterations were made to the protein or DNA in the dimer models.

### Bioinformatic Analysis of HT-SELEX, CUT&RUN, and DNA Shape Data

To compare the enrichment of dimer sites consisting of spacers with variable G/C and A/T content in HT-SELEX data, we first utilized a published position weight matrix (PWM) that was generated from the fourth cycle of a GSX2 HT-SELEX experiment^19^. We then modified this PWM to model an A/T rich and G/C rich spacer by setting equal weights of A and T or G and C respectively in the 5 positions of the spacer. The PWM of these regions are shown boxed in Figure 5A. The percentage of sequences containing each motif was determined using the known Motifs tool in HOMER^45^.

To assess the prevalence of A/T versus G/C spacer content using *in vivo* Gsx2 CUT&RUN binding data, we compared the G/C content of the spacers found in genomic dimer sites. We first identified the top quartile of dimer sites based on reads per million that were previously found in Gsx2 CUT&RUN in the mouse forebrain^18^. The dimer sites were then aligned and oriented through the known Motifs tool in HOMER. Once aligned, the spacer G/C content was calculated via the Biostrings package in R (Pagès, H., Aboyoun, R., Gentleman, R. & DebRoy, S. Biostrings: Efficient manipulation of biological strings. R package version 2.66.0.). The modeled normal distribution curve follows a mean of 0.5 and a standard deviation of 0.33. The DNA feature predictions of the six EMSA probes (sequences can be found in Table S2) were generated with DNAShapeR in R^46^.

### Electrophoretic Mobility Shift Assay (EMSA)

EMSA probes were prepared as previously described^47^. Sequences used for EMSA probes can be found in Table S2. EMSA binding reactions using Gsx2^HD^ were prepared as described previously^48^ and incubated at room temperature for 15 minutes before being run on a 7.5% polyacrylamide gel for 3 hours at 150V. EMSA binding reactions using Gsx2^167–305^ were run on a 4.5% polyacrylamide gel for 2 hours. A Li-Cor Odyssey CLx scanner was used to image all gels, and Li-Cor image studio software was used to quantify all bands. Tau factor calculations were conducted as previously described to calculate relative levels of cooperativity for each protein construct^30^. The Tau factor calculation is shown below:

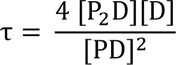

The Tau factor equation is essentially the dissociation constant of the second binding event (K_d2_) divided by the dissociation constant of the first binding event (K_d1_). This can be rewritten into individual components of dimer [P_2_D], monomer [PD], and unbound probe [D], as illustrated in the above equation. To normalize for background signal in the monomer and dimer bands, the [PD] and [P_2_D] from the empty probe lane were subtracted from the corresponding signal in the other lanes. The final equation can be given as follows:

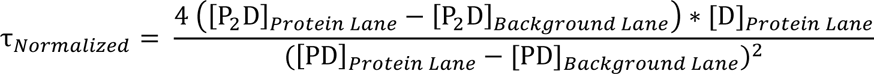

All values given in figures are the tau_Normalized_, unless otherwise noted.

**Figure S1. AlphaFold predicts full-length Gsx2 to be mostly unstructured except for the HD.** Cartoon image of AlphaFold’s structural prediction of full-length Gsx2. The structure is colored on a spectrum from red to blue representing the pLDDT value of each residue. The higher the pLDDT, the higher the accuracy of the model. Any residue with a pLDDT > 90 is expected to be modeled with high accuracy, while a pLDDT < 50 is a strong predictor of disorder^21^.

**Figure S2. The asymmetric unit of Gsx2^HD^-DNA crystals contain two complexes with a high degree of structural similarity.** (A) The asymmetric unit contains two Gsx2^HD^-DNA complexes. One Gsx2^HD^ molecule is shown in cyan, while the other is in green. DNA for both complexes is grey. (B) An alignment of all 1071 atoms from one Gsx2^HD^-DNA complex to the other complex gives a resulting RMSD value of 0.265Å. One Gsx2^HD^-DNA complex is cyan, while the other is green.

**Figure S3. Bioinformatic analysis reveals Gsx2 strongly prefers DNA dimer sites with flexible A/T-rich spacer sequences.** (A) Tau factor calculations of each sequence used in Figure 5 EMSAs show that all A/T-rich spacer sequences promote increased cooperativity compared to G/C-rich spacers. Significance was determined by one-way ANOVA with Tukey’s multiple comparisons test. (B) Measurements of the free probe from EMSAs in Figure 5 reveal no change in Gsx2^167–305^’s ability to bind the individual DNA sites regardless of the spacer sequence. Bioinformatic DNA shape analyses of these various DNA dimer site sequences showed a narrower minor groove for A/T-rich spacers (C) and increased magnitude of propellor twisting (D) and roll (F) for A/T-rich sequences, all of which are consistent with A/T-rich sequences having intrinsically more flexibility. No strong discernible pattern was observed with overall helical twist between A/T-rich and G/C-rich spacers (E).

**Figure S4. Mutating residues within the modeled Gsx2-Gsx2 binding interface diminish Gsx2’s cooperative dimerization on DNA.** (A) EMSA comparing WT^HD^ binding on the 7bpS DNA dimer site with three HD mutant constructs; S212E^HD^, L216E^HD^, and L231E^HD^. All mutants decrease cooperativity while maintaining the ability to bind DNA. (B) EMSA comparing WT^HD^ with the same three mutant constructs on the 8bpS DNA dimer site. All constructs bind equally well, with little to no cooperative dimerization observe^W^d.^T^Protein concentrations used were 0, 25, 100, and 400 nM.

**Figure S5. Triplicate EMSAs comparing WT^HD^ to I234E^HD^ show high reproducibility.** EMSA replicates comparing WT^HD^/I234E^HD^ (A-C) and WT^167–305^/I234E^167–305^ (D-F) on both the 7bpS and 8bpS DNA probes. The similarity of I234E binding on the cooperative 7bpS and non-cooperative 8bpS probes demonstrates the significant disruption to Gsx2’s ability to dimerize cooperatively on DNA. Protein concentrations of 0, 25, 100, and 400nM were used.

**Figure S6. Isothermal titration calorimetry data of Gsx2 203-264 I234E show nearly identical binding characteristics as observed with wildtype Gsx2 203-264.** (A) Isotherm of Gsx2 203-264 I234E binding to the 15mer consensus monomer site DNA shows proper stoichiometry with low nanomolar affinity, consistent with wildtype Gsx2 203-264 binding to the same 15mer consensus monomer site DNA.

