## Supplemental Data for "Cooperative Gsx2-DNA Binding Requires DNA Bending and a Novel Gsx2 Homeodomain Interface"

**Figure S1. AlphaFold predicts full-length Gsx2 to be mostly unstructured except for the HD.** Cartoon image of AlphaFold's structural prediction of full-length Gsx2. The structure is colored on a spectrum from red to blue representing the pLDDT value of each residue. The higher the pLDDT, the higher the accuracy of the model. Any residue with a pLDDT > 90 is expected to be modeled with high accuracy, while a pLDDT < 50 is a strong predictor of disorder[21].

**Figure S2. The asymmetric unit of Gsx2$^{HD}$-DNA crystals contain two complexes with a high degree of structural similarity.** (A) The asymmetric unit contains two Gsx2$^{HD}$-DNA complexes. One Gsx2$^{HD}$ molecule is shown in cyan, while the other is in green. DNA for both complexes is grey. (B) An alignment of all 1071 atoms from one Gsx2$^{HD}$-DNA complex to the other complex gives a resulting RMSD value of 0.265Å. One Gsx2$^{HD}$-DNA complex is cyan, while the other is green.

**Figure S3. Bioinformatic analysis reveals Gsx2 strongly prefers DNA dimer sites with flexible A/T-rich spacer sequences.** (A) Tau factor calculations of each sequence used in Figure 5 EMSAs show that all A/T-rich spacer sequences promote increased cooperativity compared to G/C-rich spacers. Significance was determined by one-way ANOVA with Tukey's multiple comparisons test. (B) Measurements of the free probe from EMSAs in Figure 5 reveal no change in Gsx2$^{167-305}$'s ability to bind the individual DNA sites regardless of the spacer sequence. Bioinformatic DNA shape analyses of these various DNA dimer site sequences showed a narrower minor groove for A/T-rich spacers (C) and increased magnitude of propellor twisting (D) and roll (F) for A/T-rich sequences, all of which are consistent with A/T-rich sequences having intrinsically more flexibility. No strong discernible pattern was observed with overall helical twist between A/T-rich and G/C-rich spacers (E).

**Figure S4. Mutating residues within the modeled Gsx2-Gsx2 binding interface diminish Gsx2's cooperative dimerization on DNA.** (A) EMSA comparing $WT^{HD}$ binding on the 7bpS DNA dimer site with three HD mutant constructs; $S212E^{HD}$, $L216E^{HD}$, and $L231E^{HD}$. All mutants decrease cooperativity while maintaining the ability to bind DNA. (B) EMSA comparing $WT^{HD}$ with the same three mutant constructs on the 8bpS DNA dimer site. All constructs bind equally well, with little to no cooperative dimerization observed. Protein concentrations used were 0, 25, 100, and 400 nM.

**Figure S5. Triplicate EMSAs comparing $WT^{HD}$ to $I234E^{HD}$ show high reproducibility.** EMSA replicates comparing $WT^{HD}$/$I234E^{HD}$ (A-C) and $WT^{167\text{-}305}$/$I234E^{167\text{-}305}$ (D-F) on both the 7bpS and 8bpS DNA probes. The similarity of I234E binding on the cooperative 7bpS and non-cooperative 8bpS probes demonstrates the significant disruption to Gsx2's ability to dimerize cooperatively on DNA. Protein concentrations of 0, 25, 100, and 400nM were used.

**Figure S6. Isothermal titration calorimetry data of Gsx2 203-264 I234E show nearly identical binding characteristics as observed with wildtype Gsx2 203-264.** (A) Isotherm of Gsx2 203-264 I234E binding to the 15mer consensus monomer site DNA shows proper stoichiometry with low nanomolar affinity, consistent with wildtype Gsx2 203-264 binding to the same 15mer consensus monomer site DNA.

**Table S1. Curves+ analysis of our solved Gsx2-DNA structure.**

```
 *************************************          **************
 ****   CURVES+ Version 3.0nc 09/2016   ***      *   28 Jun 22   *
 *************************************          **************

 FILE : 8EML.pdb                ftop :
 LIS  : output                  LIB  : standard
 ibld :                         sol  :
 BACK : P

 WBACK :  2.90  WBASE :  3.50  RVFAC :  7.50

 isym  :    1  itst  :    0  itnd  :    0  itdel :    1  itbkt :    0
 NAXLIM:    1

 CIRC  :    F  LINE  :    F  ZAXE  :    F  FIT   :    T  test  :    F
 ions  :    F  REFO  :    F  axfrm :    F  frames:    F


 LS fitting of standard bases ...RMS max =  0.039

 Strands =    2 Atoms =   574 Units =    28

 Combined strands have   14 levels ...

 Strand  1 has  14 bases (5'-3'): GAGCTAATTAAAGC
 Strand  2 has  14 bases (3'-5'): CTCGATTAATTTCG

 (A) BP-Axis        Xdisp   Ydisp   Inclin    Tip  Ax-bend

   1) G    2-C 15    0.14   -1.73     8.4     9.5     ---
   2) A    3-T 14   -0.52    0.15     9.5     1.5     2.0
   3) G    4-C 13   -0.11   -0.07     9.9     3.7     2.3
   4) C    5-G 12   -0.09   -0.81     4.6    -0.7     2.2
   5) T    6-A 11   -0.99   -0.87    11.4    -1.1     1.9
   6) A    7-T 10   -0.93    0.26     7.4    -1.0     2.2
   7) A    8-T  9   -0.76    0.16     2.5    -3.2     1.7
   8) T    9-A  8    0.13   -0.84    -3.8    -3.4     1.7
   9) T   10-A  7   -0.90   -1.01    -0.2    -2.7     1.9
  10) A   11-T  6    0.42   -0.00     6.2     1.9     2.0
  11) A   12-T  5   -0.70   -0.01     0.8     1.9     1.8
  12) A   13-T  4   -0.71   -0.23     0.3    -0.7     1.5
  13) G   14-C  3   -0.73   -0.25     4.7     3.9     1.8
  14) C   15-G  2   -1.00   -1.30     8.2     1.8     2.1

     Average:       -0.48   -0.47     5.0     0.8  Total bend =    19.9 (  2 to  13)

 (B) Intra-BP parameters

 Strands 1-2       Shear  Stretch Stagger  Buckle  Propel Opening

   1) G   2-C  15   -0.66   -0.51   -0.80    -1.9   -14.4    10.2
   2) A   3-T  14    0.30    0.05   -0.03    13.9   -16.4     1.3
```

| | | | | | | |
|---|---|---|---|---|---|---|
| 3) G 4-C 13 | 0.99 | -0.24 | 0.04 | 8.0 | -8.3 | -1.3 |
| 4) C 5-G 12 | -0.38 | -0.14 | 0.61 | -6.2 | -11.7 | 3.3 |
| 5) T 6-A 11 | -0.43 | 0.21 | -0.18 | 1.9 | -16.1 | 3.6 |
| 6) A 7-T 10 | 0.40 | -0.44 | -0.27 | -2.6 | -8.9 | 5.3 |
| 7) A 8-T 9 | 0.29 | -0.15 | -0.08 | 6.2 | -15.9 | 1.3 |
| 8) T 9-A 8 | -0.16 | 0.00 | 0.23 | 2.7 | -10.8 | 9.8 |
| 9) T 10-A 7 | -0.06 | 0.04 | -0.29 | 7.3 | -9.3 | 6.7 |
| 10) A 11-T 6 | 0.17 | -0.33 | -0.24 | 2.1 | -4.0 | 8.6 |
| 11) A 12-T 5 | -0.08 | -0.16 | 0.30 | 2.4 | -13.3 | -1.9 |
| 12) A 13-T 4 | -0.23 | -0.33 | 0.19 | -1.9 | -5.9 | 1.0 |
| 13) G 14-C 3 | 0.16 | 0.07 | 0.14 | 8.8 | -5.9 | -2.3 |
| 14) C 15-G 2 | -0.24 | -0.06 | -0.28 | 12.1 | -7.0 | -1.9 |
| Average: | 0.01 | -0.14 | -0.05 | 3.8 | -10.6 | 3.1 |

(C) Inter-BP

| Inter-BP | Shift | Slide | Rise | Tilt | Roll | Twist | H-Ris | H-Twi |
|---|---|---|---|---|---|---|---|---|
| 1) G 2/A 3 | 0.04 | 1.91 | 2.82 | -4.7 | -0.8 | 40.3 | 3.05 | 40.6 |
| 2) A 3/G 4 | 0.21 | -0.29 | 3.46 | -2.3 | 10.0 | 33.1 | 3.34 | 34.2 |
| 3) G 4/C 5 | 0.02 | -0.68 | 3.56 | -5.9 | 1.6 | 29.4 | 3.44 | 29.7 |
| 4) C 5/T 6 | -0.41 | -0.31 | 2.98 | 8.1 | 6.0 | 33.8 | 2.90 | 34.4 |
| 5) T 6/A 7 | 0.01 | 0.56 | 3.54 | -1.0 | 7.9 | 40.8 | 3.57 | 41.5 |
| 6) A 7/A 8 | -0.24 | -0.22 | 3.08 | -1.9 | 0.8 | 31.9 | 3.02 | 31.9 |
| 7) A 8/T 9 | 1.12 | -0.89 | 3.36 | -2.9 | -0.9 | 27.6 | 3.44 | 27.6 |
| 8) T 9/T 10 | -0.18 | -0.34 | 3.08 | 6.8 | -1.8 | 35.5 | 3.11 | 35.7 |
| 9) T 10/A 11 | 1.59 | 0.99 | 3.46 | 6.7 | 4.5 | 38.5 | 3.51 | 38.6 |
| 10) A 11/A 12 | -1.31 | 0.21 | 3.34 | -7.0 | 0.4 | 34.1 | 3.38 | 34.2 |
| 11) A 12/A 13 | -0.03 | -0.42 | 3.32 | -2.2 | -2.9 | 38.0 | 3.32 | 38.0 |
| 12) A 13/G 14 | 0.23 | -0.07 | 3.24 | 1.7 | 6.6 | 33.9 | 3.21 | 34.1 |
| 13) G 14/C 15 | 0.37 | -1.02 | 3.19 | 1.0 | 3.0 | 29.0 | 2.99 | 29.3 |
| Average: | 0.11 | -0.04 | 3.26 | -0.3 | 2.6 | 34.3 | 3.25 | 34.6 |

(D) Backbone Parameters

| Strand 1 | Alpha | Beta | Gamma | Delta | Epsil | Zeta | Chi | Phase | Ampli | Puckr |
|---|---|---|---|---|---|---|---|---|---|---|
| 1) G 2 | ---- | 90.4 | 46.8 | 139.1 | -149.5 | -166.0 | -64.3 | 158.0 | 38.2 | C2'en |
| 2) A 3 | -44.8 | 117.1 | 67.5 | 117.9 | -171.9 | -116.9 | -110.4 | 126.2 | 33.6 | C1'ex |
| 3) G 4 | 4.1 | 144.6 | -0.5 | 143.6 | -140.3 | -145.7 | -99.7 | 152.9 | 44.5 | C2'en |
| 4) C 5 | -63.3 | 131.5 | 58.7 | 79.2 | -174.9 | -74.2 | -160.6 | 53.9 | 44.1 | C4'ex |
| 5) T 6 | -71.3 | -174.1 | 66.8 | 140.8 | -166.4 | -120.8 | -113.6 | 161.3 | 34.5 | C2'en |
| 6) A 7 | 39.0 | 172.4 | -52.3 | 156.2 | 179.8 | -100.7 | -99.7 | -157.8 | 35.5 | C3'ex |
| 7) A 8 | -42.2 | 175.4 | 29.9 | 141.5 | -155.4 | -122.9 | -96.6 | 160.5 | 39.8 | C2'en |
| 8) T 9 | 62.6 | -129.3 | -112.2 | 124.0 | -160.2 | -80.5 | -147.1 | 145.5 | 16.1 | C2'en |
| 9) T 10 | -70.2 | 176.6 | 56.6 | 126.9 | -166.2 | -93.5 | -108.7 | 137.6 | 35.8 | C1'ex |
| 10) A 11 | -58.6 | 167.9 | 46.3 | 140.2 | -97.2 | 166.3 | -95.9 | 144.2 | 48.4 | C2'en |
| 11) A 12 | -74.3 | 136.3 | 46.1 | 149.1 | -129.8 | -136.1 | -111.1 | 157.1 | 46.7 | C2'en |
| 12) A 13 | -64.1 | 149.2 | 34.6 | 102.9 | -177.7 | -93.1 | -131.1 | 95.8 | 35.0 | O1'en |
| 13) G 14 | -52.4 | 171.3 | 50.7 | 137.3 | -168.9 | -106.6 | -106.5 | 153.3 | 37.1 | C2'en |
| 14) C 15 | -69.4 | 156.0 | 80.2 | 113.2 | ---- | ---- | -138.0 | 116.8 | 47.7 | C1'ex |

| Strand 2 | Alpha | Beta | Gamma | Delta | Epsil | Zeta | Chi | Phase | Ampli | Puckr |
|---|---|---|---|---|---|---|---|---|---|---|
| 1) C 15 | -17.9 | 155.9 | 37.1 | 146.6 | ---- | ---- | -88.6 | -173.1 | 33.0 | C3'ex |

| | | | | | | | | | | | | |
|---|---|---|---|---|---|---|---|---|---|---|---|---|
| 2) | T | 14 | 7.5 | 138.3 | 3.6 | 124.7 | -177.8 | -133.7 | -111.0 | 133.8 | 36.7 | C1'ex |
| 3) | C | 13 | -40.6 | 158.6 | 47.7 | 127.9 | -163.7 | -133.9 | -126.2 | 136.4 | 40.0 | C1'ex |
| 4) | G | 12 | -49.0 | 164.9 | 40.1 | 123.5 | -173.3 | -118.9 | -117.4 | 129.8 | 37.2 | C1'ex |
| 5) | A | 11 | -0.1 | 143.4 | 13.4 | 150.5 | -173.6 | -117.2 | -89.1 | 169.6 | 41.8 | C2'en |
| 6) | T | 10 | 19.4 | 155.9 | -14.9 | 150.9 | -165.0 | -149.7 | -92.8 | 170.5 | 42.4 | C2'en |
| 7) | T | 9 | -64.2 | 176.9 | 57.2 | 118.6 | -175.5 | -116.9 | -126.6 | 126.0 | 36.4 | C1'ex |
| 8) | A | 8 | -67.1 | 179.0 | 55.1 | 111.7 | 178.9 | -91.3 | -127.6 | 111.9 | 32.5 | C1'ex |
| 9) | A | 7 | -55.9 | 147.9 | 38.1 | 141.3 | 170.1 | -91.3 | -105.7 | 175.0 | 32.1 | C2'en |
| 10) | T | 6 | -59.4 | -178.9 | 43.3 | 141.6 | -121.9 | -179.3 | -99.5 | 148.3 | 45.7 | C2'en |
| 11) | T | 5 | -60.8 | 167.2 | 56.7 | 120.9 | -179.6 | -85.0 | -121.4 | 128.2 | 38.2 | C1'ex |
| 12) | T | 4 | -64.3 | 172.8 | 62.8 | 131.7 | -171.2 | -104.3 | -115.2 | 152.5 | 32.9 | C2'en |
| 13) | C | 3 | -55.6 | -177.3 | 45.5 | 136.5 | -164.8 | -109.8 | -128.3 | 152.1 | 35.0 | C2'en |
| 14) | G | 2 | ---- | 125.6 | -2.4 | 95.4 | -175.0 | -75.5 | -159.1 | 53.9 | 30.7 | C4'ex |

(E) Groove parameters

| Level | | W12 | D12 | W21 | D21 |
|---|---|---|---|---|---|
| 1.5 | | | | | |
| 2.0 | A 3 | | | | |
| 2.5 | | | | | |
| 3.0 | G 4 | 8.1 | 4.1 | | |
| 3.5 | | 7.5 | 4.8 | | |
| 4.0 | C 5 | 6.8 | 5.0 | | |
| 4.5 | | 7.1 | 4.8 | 10.8 | 6.5 |
| 5.0 | T 6 | 7.2 | 4.4 | 10.4 | 6.9 |
| 5.5 | | 6.9 | 4.7 | 10.6 | 6.9 |
| 6.0 | A 7 | 6.4 | 5.0 | 11.0 | 6.5 |
| 6.5 | | 6.2 | 4.8 | 10.6 | 6.2 |
| 7.0 | A 8 | 5.9 | 4.5 | 10.4 | 5.8 |
| 7.5 | | 5.5 | 5.1 | 11.5 | 5.7 |
| 8.0 | T 9 | 5.2 | 5.3 | 12.8 | 5.0 |
| 8.5 | | 5.5 | 5.0 | 13.3 | 5.3 |
| 9.0 | T 10 | 5.8 | 4.6 | 13.4 | 5.7 |
| 9.5 | | 4.6 | 5.9 | 13.5 | 5.1 |
| 10.0 | A 11 | 3.8 | 6.9 | 12.9 | 3.9 |
| 10.5 | | 4.7 | 6.1 | 12.2 | 4.2 |
| 11.0 | A 12 | 6.0 | 5.0 | | |
| 11.5 | | 6.8 | 4.9 | | |
| 12.0 | A 13 | | | | |
| 12.5 | | | | | |
| 13.0 | G 14 | | | | |
| 13.5 | | | | | |

**Table S2. Oligonucleotides used for EMSAs, ITC, X-ray crystallography, and site-directed mutagenesis.**

| **EMSA** | |
|---|---|
| AAATT Forward | 5'-TCCAAC**<u>TAAT</u>**TAAAATT**<u>TAAT</u>**TCG*TAGTGCGGGCGTGGCT*-3' |
| AAATT Reverse | 5'-CGA**<u>ATTA</u>**AATTTTA**<u>ATTA</u>**GTTGGA-3' |
| AATTT Forward | 5'-TCCAAC**<u>TAAT</u>**TAAATTT**<u>TAAT</u>**TCG*TAGTGCGGGCGTGGCT*-3' |
| AATTT Reverse | 5'-CGA**<u>ATTA</u>**AAATTTA**<u>ATTA</u>**GTTGGA-3' |
| ATATT Forward | 5'-TCCAAC**<u>TAAT</u>**TAATATT**<u>TAAT</u>**TCG*TAGTGCGGGCGTGGCT*-3' |
| ATATT Reverse | 5'-CGA**<u>ATTA</u>**AATATTA**<u>ATTA</u>**GTTGGA-3' |
| GCGGG Forward | 5'-TCCAAC**<u>TAAT</u>**TAGCGGG**<u>TAAT</u>**TCG*TAGTGCGGGCGTGGCT*-3' |
| GCGGG Reverse | 5'-CGA**<u>ATTA</u>**CCCGCTA**<u>ATTA</u>**GTTGGA-3' |
| GCCGG Forward | 5'-TCCAAC**<u>TAAT</u>**TAGCCGG**<u>TAAT</u>**TCG*TAGTGCGGGCGTGGCT*-3' |
| GCCGG Reverse | 5'-CGA**<u>ATTA</u>**CCGGCTA**<u>ATTA</u>**GTTGGA-3' |
| GGGCC Forward | 5'-TCCAAC**<u>TAAT</u>**TAGGGCC**<u>TAAT</u>**TCG*TAGTGCGGGCGTGGCT*-3' |
| GGGCC Reverse | 5'-CGA**<u>ATTA</u>**GGCCCTA**<u>ATTA</u>**GTTGGA-3' |
| Gsx2 7bpS Forward | 5'-TCCAAC**<u>TAAT</u>**TAAAATT**<u>TAAT</u>**TCG*TAGTGCGGGCGTGGCT*-3' |
| Gsx2 7bpS Reverse | 5'-CGA**<u>ATTA</u>**AATTTTA**<u>ATTA</u>**GTTGGA-3' |
| Gsx2 8bpS Forward | 5'-TCCAAC**<u>TAAT</u>**TAAAGATT**<u>TAAT</u>**TCG*TAGTGCGGGCGTGGCT*-3' |
| Gsx2 8bpS Reverse | 5'-CGA**<u>ATTA</u>**AATCTTTA**<u>ATTA</u>**GTTGGA-3' |
| Linker for Probe | 5'-IRDye700-*AGCCACGCCCGCACTA*-3' |

| **Isothermal Titration Calorimetry** | |
|---|---|
| Consensus Site Forward | 5'-TGAGC**<u>TAATTA</u>**AAGC-3' |
| Consensus Site Reverse | 5'-CTCG**<u>ATTAAT</u>**TTCGA-3' |
| Common Q50 Site Forward | 5'-TGAGC**<u>TAATGG</u>**AAGC-3' |
| Common Q50 Site Reverse | 5'-CTCG**<u>ATTACC</u>**TTCGA-3' |

| **X-ray Crystallography** | |
|---|---|
| Consensus Site Forward | 5'-TGAGC**<u>TAATTA</u>**AAGC-3' |
| Consensus Site Reverse | 5'-CTCG**<u>ATTAAT</u>**TTCGA-3' |

| **Site-Directed Mutagenesis** | |
|---|---|
| Gsx2 S212E Forward | 5'-GAGGACAGCGTTTACCGAGACGCAGCTCCTGGAGC-3' |
| Gsx2 S212E Reverse | 5'-GCTCCAGGAGCTGCGTCTCGGTAAACGCTGTCCTC-3' |
| Gsx2 L216E Forward | 5'-CAGCACGCAGCTCGAGGAGCTGGAGCGA-3' |

| | |
|---|---|
| Gsx2 L216E Reverse | 5'-TCGCTCCAGCTCCTCGAGCTGCGTGCTG-3' |
| Gsx2 L231E Forward | 5'-CCAATATGTACCTGTCCCGAGAGCGGAGAATCGAGATCGC-3' |
| Gsx2 L231E Reverse | 5'-GCGATCTCGATTCTCCGCTCTCGGGACAGGTACATATTGG-3' |
| Gsx2 I234E Forward | 5'-GTCCCGACTCCGGAGAGAGGAGATCGCGACATACC-3' |
| Gsx2 I234E Reverse | 5'-GGTATGTCGCGATCTCCTCTCTCCGGAGTCGGGAC-3' |

*Binding sites are bold and underlined. Linker sequences are italicized.

Figure S1

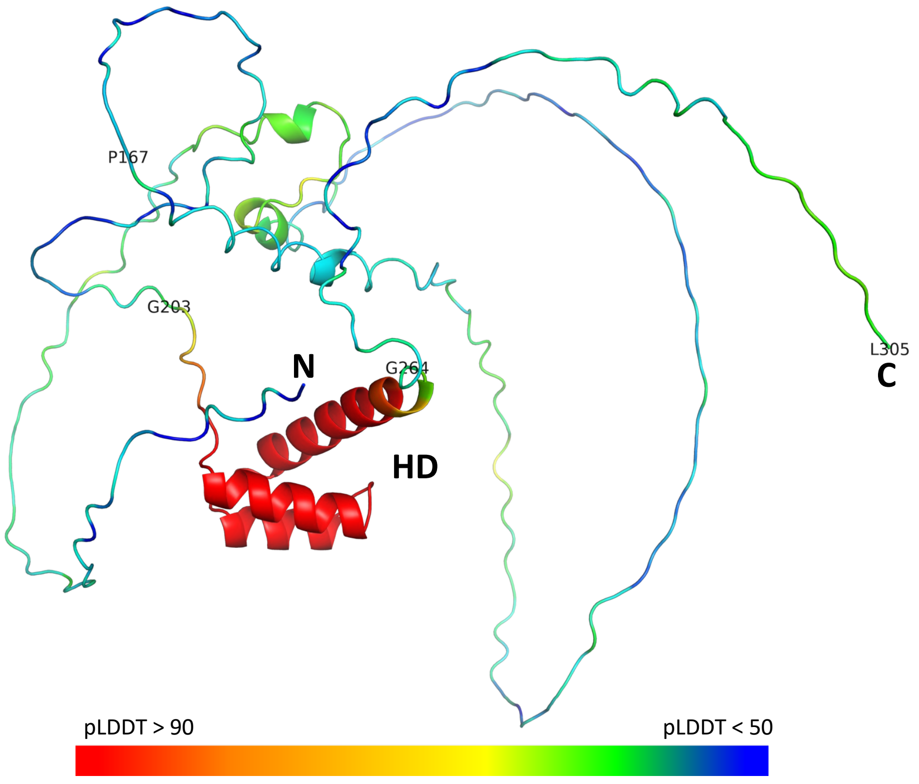

Figure S2

A

B

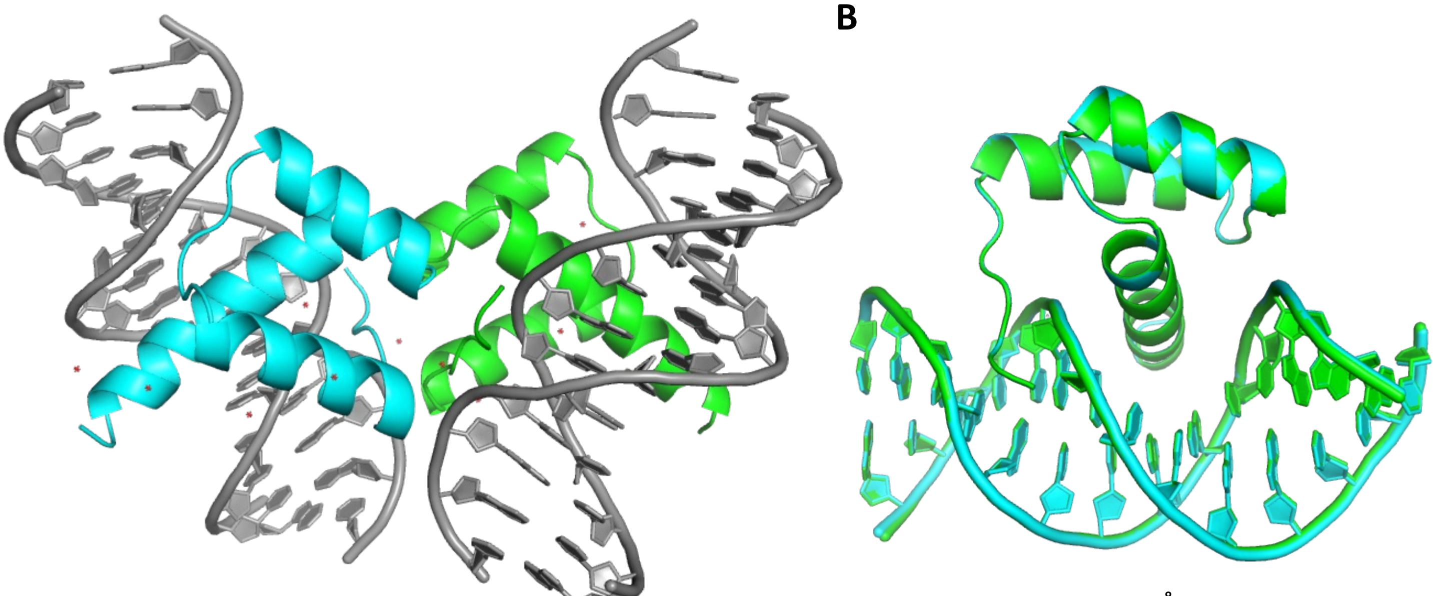

RMSD = 0.265Å (1071 atoms)

### Figure S3

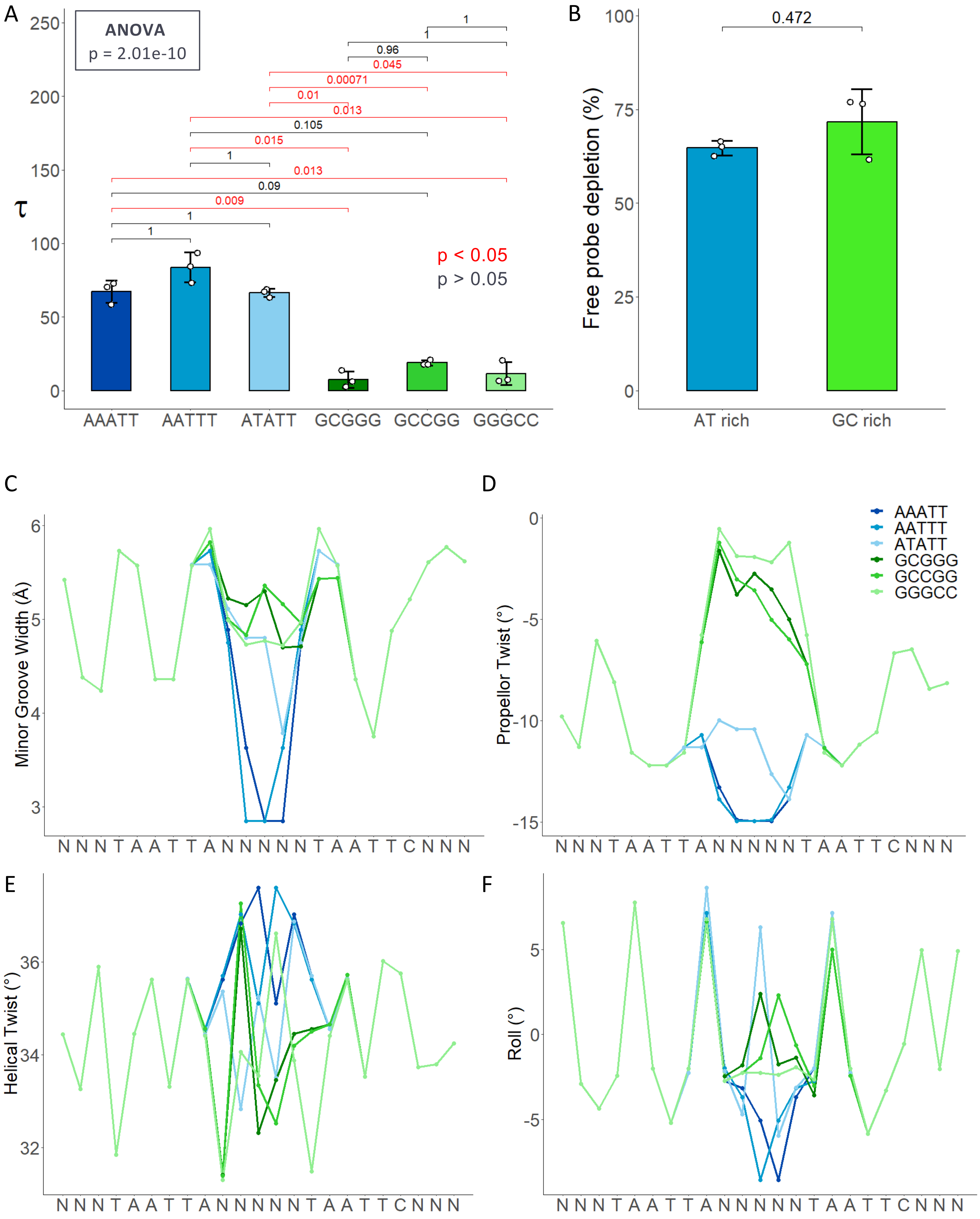
A
ANOVA
p = 2.01e-10
τ
p < 0.05
p > 0.05
AAATT
AATTT
ATATT
GCGGG
GCCGG
GGGCC
B
0.472
Free probe depletion (%)
AT rich
GC rich
C
Minor Groove Width (Å)
D
Propellor Twist (°)
E
Helical Twist (°)
F
Roll (°)
N N N T A A T T A N N N N N N T A A T T C N N N

Figure S4

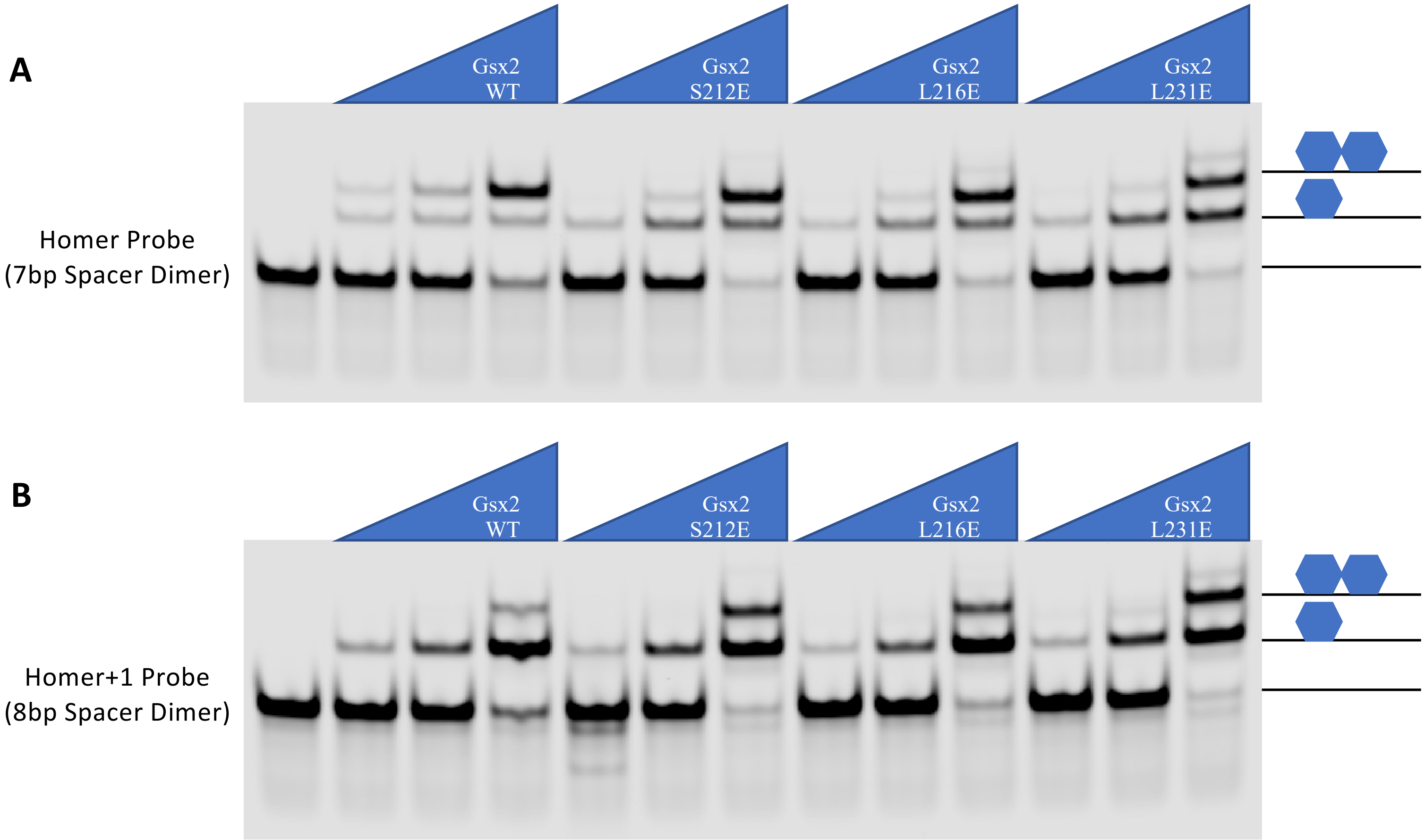
A
Gsx2
WT
Gsx2
S212E
Gsx2
L216E
Gsx2
L231E
Homer Probe
(7bp Spacer Dimer)
B
Gsx2
WT
Gsx2
S212E
Gsx2
L216E
Gsx2
L231E
Homer+1 Probe
(8bp Spacer Dimer)

Figure S5

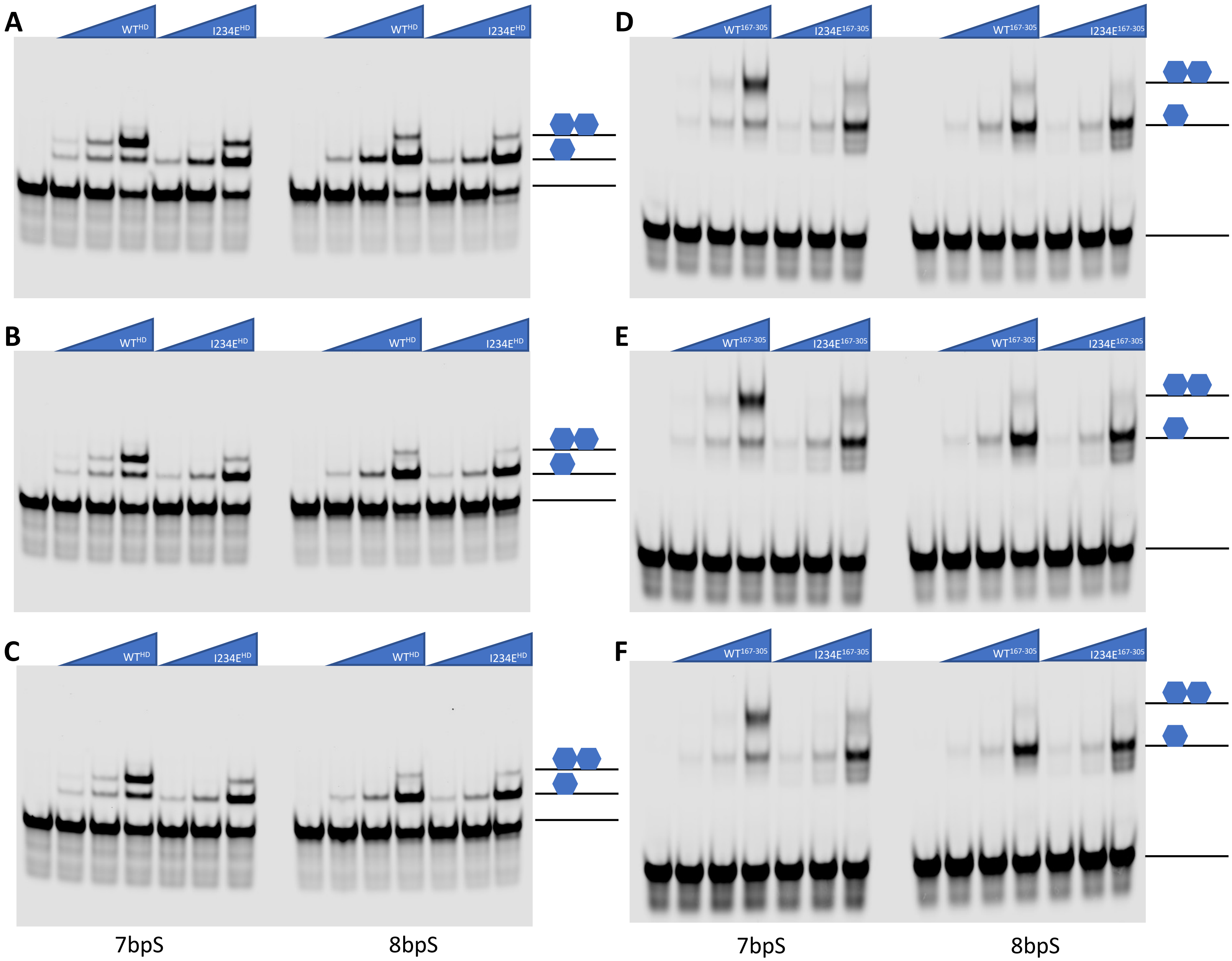

A
WT^HD
I234E^HD
WT^HD
I234E^HD
B
WT^HD
I234E^HD
WT^HD
I234E^HD
C
WT^HD
I234E^HD
WT^HD
I234E^HD
7bpS
8bpS
D
WT^167-305
I234E^167-305
WT^167-305
I234E^167-305
E
WT^167-305
I234E^167-305
WT^167-305
I234E^167-305
F
WT^167-305
I234E^167-305
WT^167-305
I234E^167-305
7bpS
8bpS

### Figure S6

#### TAATTA Motif

A

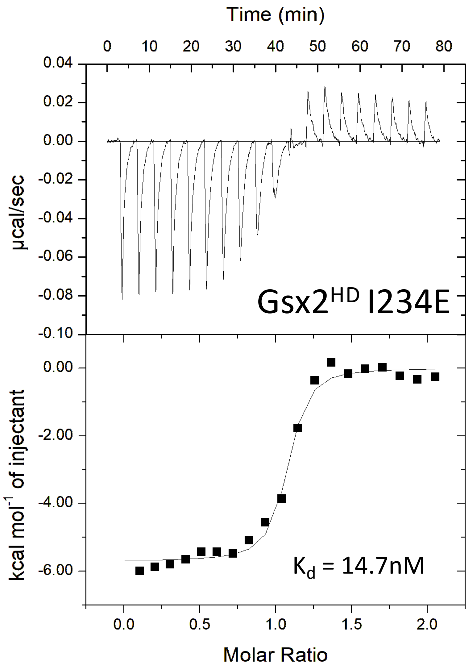

### Full wwPDB X-ray Structure Validation Report ⓘ

Oct 11, 2022 – 05:21 PM EDT

| | | |
|---|---|---|
| PDB ID | : | 8EML |
| Title | : | Crystal Structure of Gsx2 Homeodomain in Complex with DNA |
| Deposited on | : | 2022-09-28 |
| Resolution | : | 2.21 Å(reported) |

**This wwPDB validation report is for manuscript review**

This is a Full wwPDB X-ray Structure Validation Report.

This report is produced by the wwPDB biocuration pipeline after annotation of the structure.

We welcome your comments at **
A user guide is available at
https://www.wwpdb.org/validation/2017/XrayValidationReportHelp
with specific help available everywhere you see the ⓘ symbol.

The types of validation reports are described at
http://www.wwpdb.org/validation/2017/FAQs#types.

The following versions of software and data (see references ⓘ) were used in the production of this report:

| | | |
|---|---|---|
| MolProbity | : | 4.02b-467 |
| Mogul | : | 1.8.5 (274361), CSD as541be (2020) |
| Xtriage (Phenix) | : | 1.13 |
| EDS | : | 2.31.2 |
| Percentile statistics | : | 20191225.v01 (using entries in the PDB archive December 25th 2019) |
| Refmac | : | 5.8.0158 |
| CCP4 | : | 7.0.044 (Gargrove) |
| Ideal geometry (proteins) | : | Engh & Huber (2001) |
| Ideal geometry (DNA, RNA) | : | Parkinson et al. (1996) |
| Validation Pipeline (wwPDB-VP) | : | 2.31.2 |

For Manuscript Review

### 1 Overall quality at a glance (i)

The following experimental techniques were used to determine the structure:
*X-RAY DIFFRACTION*

The reported resolution of this entry is 2.21 Å.

Percentile scores (ranging between 0-100) for global validation metrics of the entry are shown in the following graphic. The table shows the number of entries on which the scores are based.

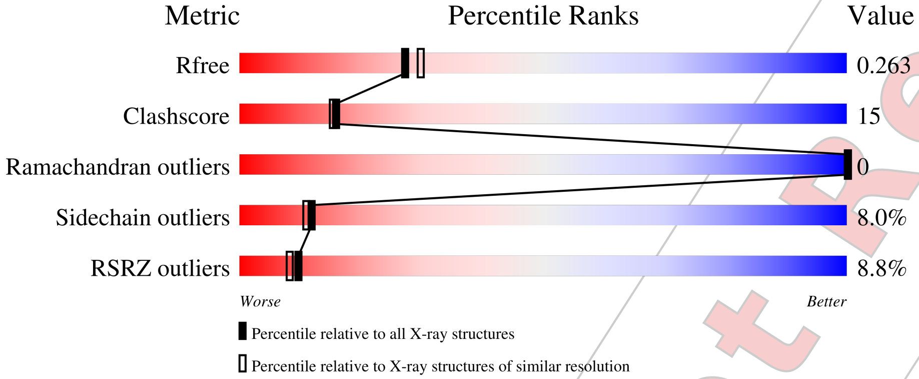


| Metric | Whole archive (#Entries) | Similar resolution (#Entries, resolution range(Å)) |
|---|---|---|
| $R_{free}$ | 130704 | 5912 (2.24-2.20) |
| Clashscore | 141614 | 6646 (2.24-2.20) |
| Ramachandran outliers | 138981 | 6543 (2.24-2.20) |
| Sidechain outliers | 138945 | 6544 (2.24-2.20) |
| RSRZ outliers | 127900 | 5797 (2.24-2.20) |

The table below summarises the geometric issues observed across the polymeric chains and their fit to the electron density. The red, orange, yellow and green segments of the lower bar indicate the fraction of residues that contain outliers for >=3, 2, 1 and 0 types of geometric quality criteria respectively. A grey segment represents the fraction of residues that are not modelled. The numeric value for each fraction is indicated below the corresponding segment, with a dot representing fractions <=5% The upper red bar (where present) indicates the fraction of residues that have poor fit to the electron density. The numeric value is given above the bar.

| Mol | Chain | Length | Quality of chain |
|---|---|---|---|
| 1 | A | 67 | 3% / 58% 24% • • 13% |
| 1 | B | 67 | 6% / 58% 22% • 15% |
| 2 | C | 15 | 7% / 33% 60% 7% |
| 2 | E | 15 | 13% / 40% 53% 7% |

*Continued on next page...*

For Manuscript Review

*Continued from previous page...*

| Mol | Chain | Length | Quality of chain |
|---|---|---|---|
| 3 | D | 15 | 20% / 60% / 33% / 7% |
| 3 | F | 15 | 20% / 67% / 27% / 7% |

The following table lists non-polymeric compounds, carbohydrate monomers and non-standard residues in protein, DNA, RNA chains that are outliers for geometric or electron-density-fit criteria:

| Mol | Type | Chain | Res | Chirality | Geometry | Clashes | Electron density |
|---|---|---|---|---|---|---|---|
| 4 | PGE | A | 301 | - | - | X | - |

For Manuscript Review

#### 2 Entry composition (i)

There are 5 unique types of molecules in this entry. The entry contains 2123 atoms, of which 0 are hydrogens and 0 are deuteriums.

In the tables below, the ZeroOcc column contains the number of atoms modelled with zero occupancy, the AltConf column contains the number of residues with at least one atom in alternate conformation and the Trace column contains the number of residues modelled with at most 2 atoms.

- Molecule 1 is a protein called GS homeobox 2.

| Mol | Chain | Residues | Atoms | | | | | ZeroOcc | AltConf | Trace |
|---|---|---|---|---|---|---|---|---|---|---|
| 1 | B | 57 | Total 473 | C 300 | N 87 | O 84 | S 2 | 0 | 0 | 0 |
| 1 | A | 58 | Total 477 | C 302 | N 88 | O 85 | S 2 | 0 | 0 | 0 |

There are 10 discrepancies between the modelled and reference sequences:

| Chain | Residue | Modelled | Actual | Comment | Reference |
|---|---|---|---|---|---|
| B | 198 | GLY | - | expression tag | UNP P31316 |
| B | 199 | PRO | - | expression tag | UNP P31316 |
| B | 200 | ALA | - | expression tag | UNP P31316 |
| B | 201 | ALA | - | expression tag | UNP P31316 |
| B | 202 | ALA | - | expression tag | UNP P31316 |
| A | 198 | GLY | - | expression tag | UNP P31316 |
| A | 199 | PRO | - | expression tag | UNP P31316 |
| A | 200 | ALA | - | expression tag | UNP P31316 |
| A | 201 | ALA | - | expression tag | UNP P31316 |
| A | 202 | ALA | - | expression tag | UNP P31316 |

- Molecule 2 is a DNA chain called DNA (5'-D(P*GP*AP*GP*CP*TP*AP*AP*TP*TP*AP *AP*AP*GP*C)-3').

| Mol | Chain | Residues | Atoms | | | | | ZeroOcc | AltConf | Trace |
|---|---|---|---|---|---|---|---|---|---|---|
| 2 | E | 14 | Total 290 | C 138 | N 57 | O 81 | P 14 | 0 | 0 | 0 |
| 2 | C | 14 | Total 290 | C 138 | N 57 | O 81 | P 14 | 0 | 0 | 0 |

- Molecule 3 is a DNA chain called DNA (5'-D(P*GP*CP*TP*TP*TP*AP*AP*TP*TP*AP *GP*CP*TP*C)-3').

For Manuscript Review

| Mol | Chain | Residues | Atoms | | | | | ZeroOcc | AltConf | Trace |
|---|---|---|---|---|---|---|---|---|---|---|
| 3 | F | 14 | Total 284 | C 137 | N 46 | O 87 | P 14 | 0 | 0 | 0 |
| 3 | D | 14 | Total 284 | C 137 | N 46 | O 87 | P 14 | 0 | 0 | 0 |

- Molecule 4 is TRIETHYLENE GLYCOL (three-letter code: PGE) (formula: $C_6H_{14}O_4$).

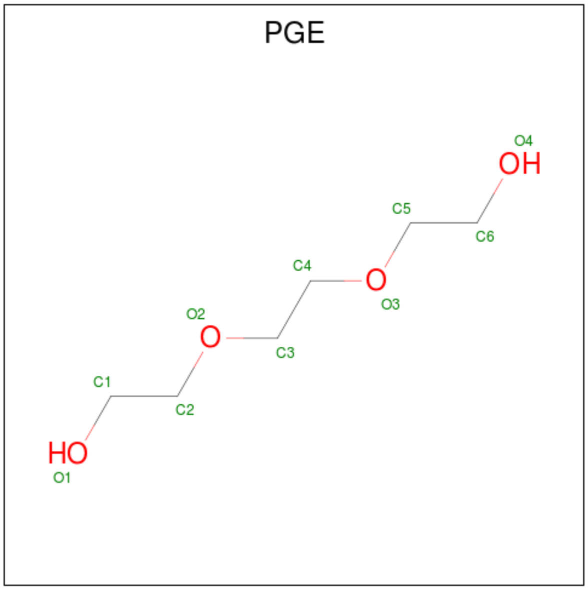


| Mol | Chain | Residues | Atoms | | | ZeroOcc | AltConf |
|---|---|---|---|---|---|---|---|
| 4 | A | 1 | Total 10 | C 6 | O 4 | 0 | 0 |

- Molecule 5 is water.

| Mol | Chain | Residues | Atoms | | ZeroOcc | AltConf |
|---|---|---|---|---|---|---|
| 5 | B | 4 | Total 4 | O 4 | 0 | 0 |
| 5 | E | 1 | Total 1 | O 1 | 0 | 0 |
| 5 | F | 3 | Total 3 | O 3 | 0 | 0 |
| 5 | A | 3 | Total 3 | O 3 | 0 | 0 |
| 5 | C | 2 | Total 2 | O 2 | 0 | 0 |
| 5 | D | 2 | Total 2 | O 2 | 0 | 0 |

For Manuscript Review

#### 3 Residue-property plots (i)

These plots are drawn for all protein, RNA, DNA and oligosaccharide chains in the entry. The first graphic for a chain summarises the proportions of the various outlier classes displayed in the second graphic. The second graphic shows the sequence view annotated by issues in geometry and electron density. Residues are color-coded according to the number of geometric quality criteria for which they contain at least one outlier: green = 0, yellow = 1, orange = 2 and red = 3 or more. A red dot above a residue indicates a poor fit to the electron density (RSRZ > 2). Stretches of 2 or more consecutive residues without any outlier are shown as a green connector. Residues present in the sample, but not in the model, are shown in grey.

- Molecule 1: GS homeobox 2

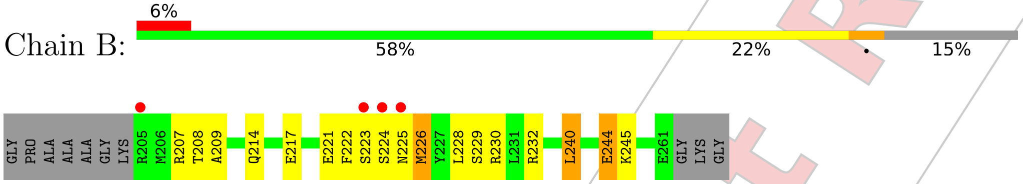


- Molecule 1: GS homeobox 2

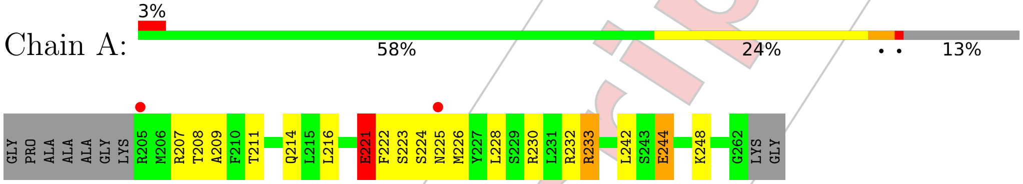


- Molecule 2: DNA (5'-D(P*GP*AP*GP*CP*TP*AP*AP*TP*TP*AP*AP*AP*GP*C)-3' )

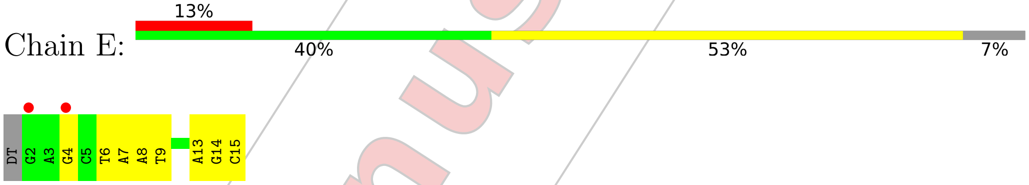


- Molecule 2: DNA (5'-D(P*GP*AP*GP*CP*TP*AP*AP*TP*TP*AP*AP*AP*GP*C)-3' )

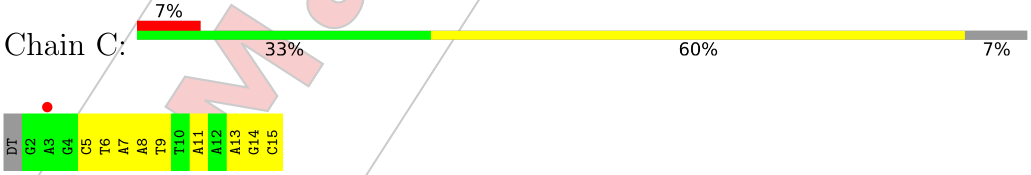


- Molecule 3: DNA (5'-D(P*GP*CP*TP*TP*TP*AP*AP*TP*TP*AP*GP*CP*TP*C)-3')

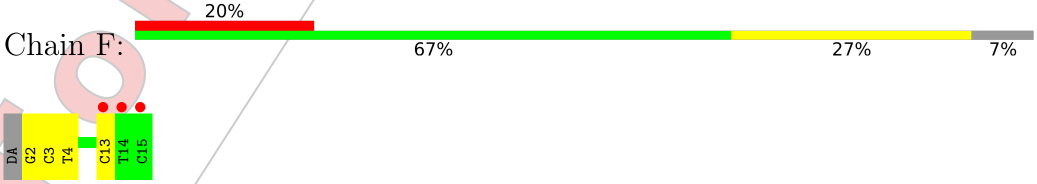


For Manuscript Review

• Molecule 3: DNA (5'-D(P*GP*CP*TP*TP*TP*AP*AP*TP*TP*AP*GP*CP*TP*C)-3')

Chain D:

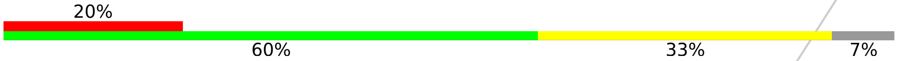


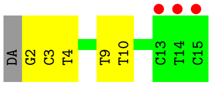


For Manuscript Review

### 4 Data and refinement statistics (i)

| Property | Value | Source |
|---|---|---|
| Space group | P 1 21 1 | Depositor |
| Cell constants<br>a, b, c, $\alpha$, $\beta$, $\gamma$ | 37.70Å 37.65Å 107.87Å<br>90.00° 93.99° 90.00° | Depositor |
| Resolution (Å) | 36.30 – 2.21<br>36.30 – 2.21 | Depositor<br>EDS |
| % Data completeness<br>(in resolution range) | 98.9 (36.30-2.21)<br>98.9 (36.30-2.21) | Depositor<br>EDS |
| $R_{merge}$ | (Not available) | Depositor |
| $R_{sym}$ | (Not available) | Depositor |
| $< I/\sigma(I) >$ [1] | 1.36 (at 2.22Å) | Xtriage |
| Refinement program | PHENIX 1.20.1_4487 | Depositor |
| R, $R_{free}$ | 0.220 , 0.265<br>0.217 , 0.263 | Depositor<br>DCC |
| $R_{free}$ test set | 1498 reflections (9.86%) | wwPDB-VP |
| Wilson B-factor (Å$^2$) | 54.8 | Xtriage |
| Anisotropy | 0.146 | Xtriage |
| Bulk solvent $k_{sol}$(e/Å$^3$), $B_{sol}$(Å$^2$) | 0.33 , 63.0 | EDS |
| L-test for twinning[2] | $< \|L\| > = 0.48$, $< L^2 > = 0.31$ | Xtriage |
| Estimated twinning fraction | No twinning to report. | Xtriage |
| $F_o$,$F_c$ correlation | 0.95 | EDS |
| Total number of atoms | 2123 | wwPDB-VP |
| Average B, all atoms (Å$^2$) | 74.0 | wwPDB-VP |

Xtriage's analysis on translational NCS is as follows: *The largest off-origin peak in the Patterson function is 8.76% of the height of the origin peak. No significant pseudotranslation is detected.*

[1] Intensities estimated from amplitudes.
[2] Theoretical values of $< |L| >$, $< L^2 >$ for acentric reflections are 0.5, 0.333 respectively for untwinned datasets, and 0.375, 0.2 for perfectly twinned datasets.

For Manuscript Review

### 5 Model quality (i)

#### 5.1 Standard geometry (i)

Bond lengths and bond angles in the following residue types are not validated in this section: PGE

The Z score for a bond length (or angle) is the number of standard deviations the observed value is removed from the expected value. A bond length (or angle) with $|Z| > 5$ is considered an outlier worth inspection. RMSZ is the root-mean-square of all Z scores of the bond lengths (or angles).

| Mol | Chain | Bond lengths | | Bond angles | |
|---|---|---|---|---|---|
| | | RMSZ | #\|Z\| >5 | RMSZ | #\|Z\| >5 |
| 1 | A | 0.73 | 0/484 | 0.95 | 3/649 (0.5%) |
| 1 | B | 0.76 | 0/480 | 1.04 | 4/644 (0.6%) |
| 2 | C | 1.04 | 0/326 | 1.16 | 4/501 (0.8%) |
| 2 | E | 1.06 | 0/326 | 1.16 | 2/501 (0.4%) |
| 3 | D | 0.98 | 0/316 | 1.10 | 0/485 |
| 3 | F | 1.01 | 0/316 | 1.13 | 0/485 |
| All | All | 0.92 | 0/2248 | 1.08 | 13/3265 (0.4%) |

Chiral center outliers are detected by calculating the chiral volume of a chiral center and verifying if the center is modelled as a planar moiety or with the opposite hand.A planarity outlier is detected by checking planarity of atoms in a peptide group, atoms in a mainchain group or atoms of a sidechain that are expected to be planar.

| Mol | Chain | #Chirality outliers | #Planarity outliers |
|---|---|---|---|
| 1 | A | 0 | 1 |

There are no bond length outliers.

All (13) bond angle outliers are listed below:

| Mol | Chain | Res | Type | Atoms | Z | Observed(°) | Ideal(°) |
|---|---|---|---|---|---|---|---|
| 1 | B | 226 | MET | CG-SD-CE | 8.11 | 113.17 | 100.20 |
| 1 | B | 244 | GLU | OE1-CD-OE2 | -7.50 | 114.30 | 123.30 |
| 1 | A | 244 | GLU | OE1-CD-OE2 | -6.87 | 115.06 | 123.30 |
| 2 | C | 14 | DG | O5'-P-OP2 | -6.33 | 100.00 | 105.70 |
| 2 | E | 14 | DG | O5'-P-OP2 | -6.30 | 100.03 | 105.70 |
| 1 | A | 221 | GLU | OE1-CD-OE2 | -6.28 | 115.77 | 123.30 |
| 2 | C | 15 | DC | C1'-O4'-C4' | -6.18 | 103.92 | 110.10 |
| 2 | E | 15 | DC | C1'-O4'-C4' | -5.95 | 104.16 | 110.10 |
| 1 | B | 226 | MET | CB-CG-SD | 5.91 | 130.12 | 112.40 |
| 1 | A | 216 | LEU | CB-CG-CD2 | -5.42 | 101.79 | 111.00 |

*Continued on next page...*

For Manuscript Review

*Continued from previous page...*

| Mol | Chain | Res | Type | Atoms | Z | Observed(°) | Ideal(°) |
|---|---|---|---|---|---|---|---|
| 2 | C | 5 | DC | O4'-C4'-C3' | -5.25 | 102.40 | 104.50 |
| 1 | B | 244 | GLU | CG-CD-OE1 | 5.16 | 128.63 | 118.30 |
| 2 | C | 11 | DA | O4'-C1'-N9 | 5.09 | 111.56 | 108.00 |

There are no chirality outliers.

All (1) planarity outliers are listed below:

| Mol | Chain | Res | Type | Group |
|---|---|---|---|---|
| 1 | A | 233 | ARG | Sidechain |

#### 5.2 Too-close contacts (i)

In the following table, the Non-H and H(model) columns list the number of non-hydrogen atoms and hydrogen atoms in the chain respectively. The H(added) column lists the number of hydrogen atoms added and optimized by MolProbity. The Clashes column lists the number of clashes within the asymmetric unit, whereas Symm-Clashes lists symmetry-related clashes.

| Mol | Chain | Non-H | H(model) | H(added) | Clashes | Symm-Clashes |
|---|---|---|---|---|---|---|
| 1 | A | 477 | 0 | 472 | 20 | 0 |
| 1 | B | 473 | 0 | 469 | 19 | 0 |
| 2 | C | 290 | 0 | 158 | 3 | 0 |
| 2 | E | 290 | 0 | 158 | 6 | 0 |
| 3 | D | 284 | 0 | 161 | 4 | 0 |
| 3 | F | 284 | 0 | 161 | 5 | 0 |
| 4 | A | 10 | 0 | 14 | 17 | 0 |
| 5 | A | 3 | 0 | 0 | 0 | 0 |
| 5 | B | 4 | 0 | 0 | 0 | 0 |
| 5 | C | 2 | 0 | 0 | 0 | 0 |
| 5 | D | 2 | 0 | 0 | 0 | 0 |
| 5 | E | 1 | 0 | 0 | 0 | 0 |
| 5 | F | 3 | 0 | 0 | 0 | 0 |
| All | All | 2123 | 0 | 1593 | 52 | 0 |

The all-atom clashscore is defined as the number of clashes found per 1000 atoms (including hydrogen atoms). The all-atom clashscore for this structure is 15.

All (52) close contacts within the same asymmetric unit are listed below, sorted by their clash magnitude.

| Atom-1 | Atom-2 | Interatomic distance (Å) | Clash overlap (Å) |
|---|---|---|---|
| 1:A:233:ARG:HH11 | 1:A:248:LYS:HG3 | 1.16 | 1.10 |

*Continued on next page...*

For Manuscript Review

*Continued from previous page...*

| Atom-1 | Atom-2 | Interatomic distance (Å) | Clash overlap (Å) |
|---|---|---|---|
| 1:A:233:ARG:NH1 | 1:A:248:LYS:HG3 | 1.75 | 1.00 |
| 1:A:221:GLU:OE2 | 1:A:232:ARG:NH2 | 2.01 | 0.94 |
| 2:C:8:DA:H2” | 2:C:9:DT:H5” | 1.64 | 0.80 |
| 1:B:209:ALA:H | 4:A:301:PGE:H52 | 1.47 | 0.79 |
| 1:A:208:THR:HA | 4:A:301:PGE:H22 | 1.65 | 0.78 |
| 2:E:8:DA:H2” | 2:E:9:DT:H5” | 1.66 | 0.78 |
| 1:A:209:ALA:H | 4:A:301:PGE:H22 | 1.56 | 0.70 |
| 1:B:208:THR:HA | 4:A:301:PGE:H52 | 1.73 | 0.68 |
| 1:B:214:GLN:HE22 | 4:A:301:PGE:H12 | 1.58 | 0.67 |
| 1:B:214:GLN:HE22 | 4:A:301:PGE:C1 | 2.09 | 0.65 |
| 3:F:3:DC:H2’ | 3:F:4:DT:H72 | 1.78 | 0.64 |
| 1:A:233:ARG:HH11 | 1:A:248:LYS:CG | 2.05 | 0.62 |
| 1:B:228:LEU:HD22 | 1:B:232:ARG:HG2 | 1.83 | 0.61 |
| 3:D:3:DC:H2’ | 3:D:4:DT:H72 | 1.83 | 0.59 |
| 1:A:214:GLN:HE22 | 4:A:301:PGE:C6 | 2.15 | 0.59 |
| 1:A:214:GLN:OE1 | 4:A:301:PGE:H5 | 2.06 | 0.56 |
| 1:B:214:GLN:NE2 | 4:A:301:PGE:H12 | 2.23 | 0.54 |
| 1:B:209:ALA:N | 4:A:301:PGE:H52 | 2.20 | 0.53 |
| 1:B:209:ALA:H | 4:A:301:PGE:C5 | 2.21 | 0.52 |
| 1:B:214:GLN:OE1 | 4:A:301:PGE:H2 | 2.10 | 0.51 |
| 1:A:228:LEU:HD22 | 1:A:232:ARG:HG2 | 1.92 | 0.51 |
| 1:B:221:GLU:HA | 1:B:221:GLU:OE1 | 2.10 | 0.51 |
| 1:B:207:ARG:HD3 | 2:E:13:DA:H5’ | 1.92 | 0.51 |
| 1:A:222:PHE:CZ | 1:A:226:MET:HG2 | 2.47 | 0.50 |
| 1:B:207:ARG:HB2 | 2:E:13:DA:H5’ | 1.94 | 0.49 |
| 1:A:242:LEU:HD23 | 4:A:301:PGE:H62 | 1.94 | 0.49 |
| 1:A:222:PHE:HA | 1:A:225:ASN:O | 2.14 | 0.48 |
| 1:B:222:PHE:HA | 1:B:225:ASN:O | 2.14 | 0.48 |
| 1:A:209:ALA:N | 4:A:301:PGE:H22 | 2.27 | 0.47 |
| 2:E:6:DT:H2” | 2:E:7:DA:H5’ | 1.96 | 0.46 |
| 1:B:222:PHE:CE1 | 1:B:226:MET:HG3 | 2.50 | 0.46 |
| 2:C:6:DT:H2” | 2:C:7:DA:H5’ | 1.96 | 0.46 |
| 3:F:2:DG:H2’ | 3:F:3:DC:C6 | 2.51 | 0.45 |
| 3:D:2:DG:H2’ | 3:D:3:DC:C6 | 2.51 | 0.45 |
| 1:B:230:ARG:O | 1:B:230:ARG:HD2 | 2.17 | 0.45 |
| 2:E:13:DA:H5” | 1:A:211:THR:HG21 | 1.98 | 0.45 |
| 1:A:209:ALA:H | 4:A:301:PGE:C2 | 2.26 | 0.44 |
| 2:E:4:DG:H1 | 3:F:13:DC:H42 | 1.64 | 0.44 |
| 1:A:207:ARG:HD3 | 2:C:13:DA:H5’ | 2.00 | 0.43 |
| 1:A:221:GLU:OE2 | 1:A:232:ARG:CZ | 2.65 | 0.43 |
| 3:F:3:DC:H2’ | 3:F:4:DT:C7 | 2.45 | 0.43 |

*Continued on next page...*

For Manuscript Review

*Continued from previous page...*

| Atom-1 | Atom-2 | Interatomic distance (Å) | Clash overlap (Å) |
|---|---|---|---|
| 1:A:208:THR:CA | 4:A:301:PGE:H22 | 2.42 | 0.43 |
| 3:D:9:DT:H2" | 3:D:10:DT:H5' | 2.01 | 0.43 |
| 1:B:222:PHE:CZ | 1:B:226:MET:HG3 | 2.54 | 0.42 |
| 3:F:2:DG:H2" | 3:F:3:DC:H5' | 2.01 | 0.42 |
| 1:B:245:LYS:HB2 | 1:B:245:LYS:HE2 | 1.76 | 0.41 |
| 1:A:230:ARG:HH11 | 1:A:230:ARG:HD3 | 1.75 | 0.41 |
| 3:D:3:DC:H2' | 3:D:4:DT:C7 | 2.47 | 0.41 |
| 1:B:217:GLU:HB2 | 1:B:240:LEU:HD11 | 2.02 | 0.40 |
| 1:B:230:ARG:NE | 1:B:244:GLU:OE2 | 2.54 | 0.40 |
| 1:A:209:ALA:O | 4:A:301:PGE:O3 | 2.38 | 0.40 |

There are no symmetry-related clashes.

#### 5.3 Torsion angles (i)

##### 5.3.1 Protein backbone (i)

In the following table, the Percentiles column shows the percent Ramachandran outliers of the chain as a percentile score with respect to all X-ray entries followed by that with respect to entries of similar resolution.

The Analysed column shows the number of residues for which the backbone conformation was analysed, and the total number of residues.

| Mol | Chain | Analysed | Favoured | Allowed | Outliers | Percentiles | |
|---|---|---|---|---|---|---|---|
| 1 | A | 56/67 (84%) | 53 (95%) | 3 (5%) | 0 | 100 | 100 |
| 1 | B | 55/67 (82%) | 53 (96%) | 2 (4%) | 0 | 100 | 100 |
| All | All | 111/134 (83%) | 106 (96%) | 5 (4%) | 0 | 100 | 100 |

There are no Ramachandran outliers to report.

##### 5.3.2 Protein sidechains (i)

In the following table, the Percentiles column shows the percent sidechain outliers of the chain as a percentile score with respect to all X-ray entries followed by that with respect to entries of similar resolution.

The Analysed column shows the number of residues for which the sidechain conformation was analysed, and the total number of residues.

For Manuscript Review

| Mol | Chain | Analysed | Rotameric | Outliers | Percentiles | |
|---|---|---|---|---|---|---|
| 1 | A | 50/58 (86%) | 46 (92%) | 4 (8%) | 12 | 11 |
| 1 | B | 50/58 (86%) | 46 (92%) | 4 (8%) | 12 | 11 |
| All | All | 100/116 (86%) | 92 (92%) | 8 (8%) | 12 | 11 |

All (8) residues with a non-rotameric sidechain are listed below:

| Mol | Chain | Res | Type |
|---|---|---|---|
| 1 | B | 223 | SER |
| 1 | B | 224 | SER |
| 1 | B | 229 | SER |
| 1 | B | 240 | LEU |
| 1 | A | 221 | GLU |
| 1 | A | 223 | SER |
| 1 | A | 224 | SER |
| 1 | A | 244 | GLU |

Sometimes sidechains can be flipped to improve hydrogen bonding and reduce clashes. All (1) such sidechains are listed below:

| Mol | Chain | Res | Type |
|---|---|---|---|
| 1 | A | 252 | GLN |

##### 5.3.3 RNA (i)

There are no RNA molecules in this entry.

#### 5.4 Non-standard residues in protein, DNA, RNA chains (i)

There are no non-standard protein/DNA/RNA residues in this entry.

#### 5.5 Carbohydrates (i)

There are no monosaccharides in this entry.

#### 5.6 Ligand geometry (i)

1 ligand is modelled in this entry.

In the following table, the Counts columns list the number of bonds (or angles) for which Mogul statistics could be retrieved, the number of bonds (or angles) that are observed in the model and

For Manuscript Review

the number of bonds (or angles) that are defined in the Chemical Component Dictionary. The Link column lists molecule types, if any, to which the group is linked. The Z score for a bond length (or angle) is the number of standard deviations the observed value is removed from the expected value. A bond length (or angle) with $|Z| > 2$ is considered an outlier worth inspection. RMSZ is the root-mean-square of all Z scores of the bond lengths (or angles).

| Mol | Type | Chain | Res | Link | Bond lengths | | | Bond angles | | |
|---|---|---|---|---|---|---|---|---|---|---|
| | | | | | Counts | RMSZ | $\#\|Z\| > 2$ | Counts | RMSZ | $\#\|Z\| > 2$ |
| 4 | PGE | A | 301 | - | 9,9,9 | 0.83 | 0 | 8,8,8 | 1.38 | 1 (12%) |

In the following table, the Chirals column lists the number of chiral outliers, the number of chiral centers analysed, the number of these observed in the model and the number defined in the Chemical Component Dictionary. Similar counts are reported in the Torsion and Rings columns. '-' means no outliers of that kind were identified.

| Mol | Type | Chain | Res | Link | Chirals | Torsions | Rings |
|---|---|---|---|---|---|---|---|
| 4 | PGE | A | 301 | - | - | 6/7/7/7 | - |

There are no bond length outliers.

All (1) bond angle outliers are listed below:

| Mol | Chain | Res | Type | Atoms | Z | Observed($^o$) | Ideal($^o$) |
|---|---|---|---|---|---|---|---|
| 4 | A | 301 | PGE | O2-C3-C4 | -2.40 | 99.58 | 110.39 |

There are no chirality outliers.

All (6) torsion outliers are listed below:

| Mol | Chain | Res | Type | Atoms |
|---|---|---|---|---|
| 4 | A | 301 | PGE | C1-C2-O2-C3 |
| 4 | A | 301 | PGE | C6-C5-O3-C4 |
| 4 | A | 301 | PGE | O1-C1-C2-O2 |
| 4 | A | 301 | PGE | O3-C5-C6-O4 |
| 4 | A | 301 | PGE | C3-C4-O3-C5 |
| 4 | A | 301 | PGE | C4-C3-O2-C2 |

There are no ring outliers.

1 monomer is involved in 17 short contacts:

| Mol | Chain | Res | Type | Clashes | Symm-Clashes |
|---|---|---|---|---|---|
| 4 | A | 301 | PGE | 17 | 0 |

For Manuscript Review

#### 5.7 Other polymers (i)

There are no such residues in this entry.

#### 5.8 Polymer linkage issues (i)

There are no chain breaks in this entry.

For Manuscript Review

### 6 Fit of model and data (i)

#### 6.1 Protein, DNA and RNA chains (i)

In the following table, the column labelled '#RSRZ> 2' contains the number (and percentage) of RSRZ outliers, followed by percent RSRZ outliers for the chain as percentile scores relative to all X-ray entries and entries of similar resolution. The OWAB column contains the minimum, median, $95^{th}$ percentile and maximum values of the occupancy-weighted average B-factor per residue. The column labelled 'Q< 0.9' lists the number of (and percentage) of residues with an average occupancy less than 0.9.

| Mol | Chain | Analysed | <RSRZ> | #RSRZ>2 | | | OWAB(Å$^2$) | Q<0.9 |
|---|---|---|---|---|---|---|---|---|
| 1 | A | 58/67 (86%) | 0.51 | 2 (3%) | 45 | 43 | 37, 61, 105, 128 | 0 |
| 1 | B | 57/67 (85%) | 0.63 | 4 (7%) | 16 | 15 | 36, 60, 99, 110 | 0 |
| 2 | C | 14/15 (93%) | 0.48 | 1 (7%) | 16 | 14 | 56, 77, 130, 144 | 0 |
| 2 | E | 14/15 (93%) | 0.56 | 2 (14%) | 2 | 2 | 56, 77, 129, 146 | 0 |
| 3 | D | 14/15 (93%) | 1.21 | 3 (21%) | 0 | 0 | 45, 75, 127, 160 | 0 |
| 3 | F | 14/15 (93%) | 0.85 | 3 (21%) | 0 | 0 | 46, 75, 128, 157 | 0 |
| All | All | 171/194 (88%) | 0.64 | 15 (8%) | 10 | 8 | 36, 68, 120, 160 | 0 |

All (15) RSRZ outliers are listed below:

| Mol | Chain | Res | Type | RSRZ |
|---|---|---|---|---|
| 3 | D | 15 | DC | 8.5 |
| 3 | F | 15 | DC | 5.2 |
| 3 | D | 13 | DC | 3.8 |
| 3 | D | 14 | DT | 3.5 |
| 2 | C | 3 | DA | 3.0 |
| 1 | A | 225 | ASN | 2.9 |
| 1 | B | 205 | ARG | 2.9 |
| 1 | B | 225 | ASN | 2.8 |
| 3 | F | 14 | DT | 2.8 |
| 2 | E | 4 | DG | 2.7 |
| 1 | B | 223 | SER | 2.6 |
| 3 | F | 13 | DC | 2.5 |
| 1 | B | 224 | SER | 2.3 |
| 1 | A | 205 | ARG | 2.2 |
| 2 | E | 2 | DG | 2.0 |

For Manuscript Review

#### 6.2 Non-standard residues in protein, DNA, RNA chains (i)

There are no non-standard protein/DNA/RNA residues in this entry.

#### 6.3 Carbohydrates (i)

There are no monosaccharides in this entry.

#### 6.4 Ligands (i)

In the following table, the Atoms column lists the number of modelled atoms in the group and the number defined in the chemical component dictionary. The B-factors column lists the minimum, median, $95^{th}$ percentile and maximum values of B factors of atoms in the group. The column labelled 'Q< 0.9' lists the number of atoms with occupancy less than 0.9.

| Mol | Type | Chain | Res | Atoms | RSCC | RSR | B-factors($Å^2$) | Q<0.9 |
|---|---|---|---|---|---|---|---|---|
| 4 | PGE | A | 301 | 10/10 | 0.84 | 0.21 | 37,45,56,58 | 0 |

#### 6.5 Other polymers (i)

There are no such residues in this entry.

For Manuscript Review
